# Chikungunya in Colombia: a description of an epidemic within the framework of a COPCORD study

**DOI:** 10.1101/389981

**Authors:** Juan C. Rueda, Ana M. Santos, Jose-Ignacio Angarita, Rodrigo B. Giraldo, Eugenia-Lucia Saldarriaga, Jesús Giovanny Ballesteros Muñoz, Elías Forero, Hugo Valencia, Francisco Somoza, Ingris Peláez-Ballestas, Mario H. Cardiel, Paula X. Pavía, John Londono

## Abstract

During 2014 and 2015 the chikungunya virus reached Colombia unleashing an epidemic that spread throughout the whole territory. Concurrently, the Colombian Rheumatology Association was conducting a Community Oriented Program for Control of Rheumatic Diseases (COPCORD) to establish rheumatic disease prevalence in the country. Chikungunya infected patients were identified within the COPCORD population. The aim of this study was to describe the demographics, clinical characteristics and disability of patients with clinical suspicion of chikungunya infection. To confirm chikungunya infection, ELISA IgM and IgG serology was performed. From the 6528-surveyed people of the COPCORD study, 548 where included in the study because of clinical suspicion of chikungunya virus infection. Of those, 295 were positive for IgG or IgM chikungunya serology with 151 patients fulfilling WHO clinical criteria for chikungunya infection (true positives). Most patients were > 45 years (57.7%), and females (69.7%). Patients with low income and low socio-economic strata had increased risk of chikungunya infection (p = 0.00; OR: 2.36, CI: 1.47-3.77 and p = 0.00; OR: 2.81, CI: 1.90-4.17 respectively). True positive patients were associated with symmetric arthritis (p = 0.00; OR: 22.49, CI: 12.71-39.80) of ankles (p = 0.00; OR: 16.06, CI: 7.57-34.08), hands (p = 0.00; OR: 16.12, CI: 8.25-39.79), feet (p = 0.00; OR: 16.35, CI: 7.41-36.05) and elbows (p = 0.00; OR: 14.00, CI: 3.03-64.70). Most patients developed mild to moderate disability (95.2 to 100%). Our study showed that poverty and low socioeconomic status are associated with increased risk of chikungunya infection. Also, we found two distinctive phenotypes of chikungunya infection; those with positive chikungunya serology and typical clinical symptoms (true positives) and those with positive serology without clinical symptoms (false negatives). Finally, a distinctive clinical picture presented by chikungunya infected patients was found which should be considered as the hallmark for diagnostic clinical criteria.

## Introduction

In 2014, the Colombian Rheumatology Association starts the endeavour of establishing the prevalence of rheumatic diseases in the country, since there was a lack of information in this area and the data was needed to improve its impact in public health. The chosen epidemiologic strategy designed to identify rheumatic diseases in Colombia was the Community Oriented Program for Control of Rheumatic Diseases (COPCORD), which has been proven effective in other Latin American countries [1–9].

COPCORD is a low-budget, low-infrastructure, community oriented programme aimed to measure and evaluate pain and disability from rheumatic disorders in developing countries [1,2,10]. It contains three stages. A first stage where population data is collected through three successive phases; a second stage that mainly identifies risk factors and educates the community and health care providers; and a third stage where preventive and control strategies are planned, executed and maintained [2,10]. As mentioned before, the first stage, which was the only one implemented so far in Colombia, was carried out through three successive phases. A house-to-house survey by a local health worker identified patients with nontraumatic musculoskeletal (MSK) symptoms (phase I); an interview based with previously validated questionnaires executed by a health-care provider captured pain and disability (phase II), and finally (phase III) a rheumatologic diagnosis was stablished through standard evaluation by a doctor (rheumatologist or fellow in rheumatology) [2].

During phase I, a chikungunya virus (CHIKV) epidemic struck the country [11]. Since CHIKV infection main complaint is MSK symptoms, the identified cases increased to numbers that previously had not been thought. In consequence, prevalence data on soft tissue rheumatism, rheumatic regional pain syndromes, appendicular regional pain syndrome, and undifferentiated inflammatory arthritis as well as disability and pain scores could be biased. Therefore, CHIKV infected patients had to be identified within the studied population.

CHIKV is an alphavirus from the *Togaviridae* family, and a member of the Semliki Forest virus antigenic complex, that together with other alphaviruses (O’nyong-nyong, Mayaro, and Ross River) causes acute arthropathy in humans [12–14]. The virion has an icosahedral capsid, enclosed by a lipid envelope with a single-stranded, positive sense, RNA genome of approximately 12 kilobases in length, which is arranged in two open reading frames (ORF) with a junction region in between [15,16]. The 5’ ORF contains code for four non-structural proteins (nsP1-4), whereas the 3’ encodes the capsid protein (C), two surface envelope glycoproteins (E1 and E2), and two small peptides designated E3 and 6k [17,18].

A mosquito bite from an infected viremic patient is where the transmission initially starts [19]. The virus replicates for a few days, before being transmitted to another person [20]. Upon mosquito bite, the virus due to cellular tropism, infects fibroblasts in the dermis and macrophages, following an incubation period of 3 to 7 days, from where it is disseminated through lymphatics and bloodstream to joint capsule, muscle, epithelial, and endothelial cells [19]. The virus replicates eliciting viremia, fever, rash, myalgia, arthralgia and arthritis [21]. At this point the acute phase is established, lasting for approximately 2 weeks and characterized by the appearance of immunoglobulin type M (IgM) (persisting for weeks up to 3 months) followed by the production of immunoglobulin type G (IgG), which will provide antiviral immunity for years [16,19,21]. After the acute phase, CHIKV infection can progress to a chronic stage where rheumatic symptoms can last for several months to years [21].

CHIKV is an arthropod-borne virus transmitted initially by the *Aedes (Ae) aegypti* and after 2006’s epidemic in La Reunion by *Ae albopictus* due to an adaptive mutation of alanine for valine in the position 226 of the E1 glycoprotein genome (A226V) [16,22]. It is believed that the infection started in a sylvatic or enzootic cycle where the virus was maintained through arboreal vectors (*Ae africanus*, and *Ae furcifer*) using non-human primates (vervet monkeys) as hosts, and later spilled over to humans living nearby forested regions [23]. From here on, transmission from human to human via *Ae aegypti* or *albopictus* is amplified in urban settings and spread by air travel establishing an epidemic cycle [23].

The first isolated cases of CHIKV were reported in July 1952 along the coastal plateaus of Mawia, Makonde and Rondo, what is known today as Tanzania [24]. The people of this region named the disease chikungunya, which translates to “the one that bends up the joints”[25]. Phylogenetic studies indicated that CHIKV originated in Africa over 500 years ago and determined that a common lineage diverged into two branches termed West African (WA) and East/Central/South African (ECSA) [12,13,26,27]. While the ECSA lineage spread outside Africa causing multiple urban epidemics in Asia almost 150 years ago, WA lineage maintained local outbreaks in Africa through enzootic transmission [19,27]. It was in Asia were the ECSA lineage kept circulating and evolving into a separate genotype called Asian lineage [27]. In the earlies 2000, the ECSA lineage reached Kenya and from there expanded to islands in the Indian Ocean, India, and Southeast Asia creating an unprecedented epidemic and again evolving into a new lineage with the aforementioned A226V mutation (Indian Ocean Lineage or IOL) [22,28–30]. This allowed the virus to use *Ae albopictus* as a new vector which has a higher altitude tolerance and therefore increasing the disease’s reach to more temperate regions like southern France and northern Italy [31–34]. During the last decade CHIKV continued to cause epidemics in the Pacific Islands, Indian subcontinent, Oceania and Southeast Asia [35–37]. Finally, in 2013 the CHIKV Asian lineage arrived to the Western Hemisphere with the first autochthonous cases reported in the Island of Saint Martin [38]. From there, the virus rapidly spread throughout the Caribbean, Central and South America, affecting 42 countries by 2015 [39].

In August 2014, CHIKV arrives to the northern region of Colombia causing an epidemic that left 106.763 reported cases in 2014 and 361.004 in 2015 and spanning throughout the whole territory (32 state departments) [11,40–43]. According to the Pan-American Health Organization (PAHO) statistics, Colombia was the third country with most cumulative cases (294.831) in the Americas between December 2013 to December 2017 behind Dominican Republic (539.362) and Brazil (773.010) [44]. At the end of 2015 the Colombian Health Ministry declares the end of the epidemic, however cases continued to be reported in the following years up to this date [45–47]. The following study describes in detail the clinical presentation and disability of a CHIKV epidemic analysed by WHO criteria, serology status, and duration of symptoms, in six representative cities of the country (Bogotá, Medellín, Cali, Barranquilla, Bucaramanga and Cúcuta), within the framework of a COPCORD study during 2015 and 2016.

## Materials and methods

### COPCORD methodology

This was a cross-sectional, analytical study designed for people over 18 years-old. A probabilistic stratified sampling method was used throughout three stages. The first sampling stage consisted of selecting cartographic areas in each city, as defined by the Colombian Statistics Administration Department (DANE, Departamento Administrativo Nacional de Estadística). The second sampling stage involved blocking each sector using an urban analysis tool which classifies cities into blocks, houses, households and people (VIHOPE). The third stage concerned the homes in each block; all household members were surveyed. Sample size was calculated at 6528 individuals for a 1.5 sampling design effect and 14% sampling error.

### Data recollection

The COPCORD questionnaire adapted for Colombia, was used during the first stage by standardised interviewers [48]. Also, at this stage, interviewers were ordered to ask about CHIKV related symptoms like fever, rash, myalgia, and fatigue as well as possible diagnosis of CHIKV infection by a health care worker or by the patient. If positive, the patients were labelled as suspected case of CHIKV infection and programmed for a later examination by a study physician. In the second examination, the physician confirmed suspicion of CHIKV infection according to WHO criteria, a specific designed questionnaire was used to gather data and blood samples were taken [49].

### Case definitions

Case definitions according to WHO criteria were as follows [49]: Clinical criteria: acute onset of fever > 38.5°C and severe arthralgia/arthritis not explained by other medical conditions.

Epidemiological criteria: residing or having visited epidemic areas, having reported transmission within 15 days prior to the onset of symptoms

Laboratory criteria: presence of virus-specific IgM antibodies in single serum sample collected in acute or convalescent stage; or four-fold increase in IgG values in samples collected at least three weeks apart (the latter definition was changed to presence of virus-specific IgG antibodies in single serum sample during any stage of the disease because our population was immunologically naïve; there are no reports of CHIKV infection prior to this epidemic).

Possible case: a patient meeting clinical criteria.

Probable case: a patient meeting both clinical and epidemiological criteria.

Confirmed case: a patient meeting laboratory criteria, irrespective of clinical presentation. EuroQol-5D (EQ-5D) was used for estimating health status and the Health Assessment Questionnaire Disability Index (HAQ-DI) for measuring functional capacity [50].

### Blood sample and Chain of custody

With the consent of the patient, proper asepsis and considering the established protocols, venous blood samples were taken by trained personnel, linked to La Universidad de La Sabana. The sample was transported to the Laboratory of Immunology of the Biomedical Campus of La Universidad de La Sabana, adhering to the protocol of chain of custody, which included a triple packaging of the sample and conservation of the cold chain. After obtaining the blood sample by venepuncture and using the vacuum system for collection in dry tubes, the blood was centrifuged at 3500 rpm for 10 minutes and the serum was separated into cryovials. The samples were stored and refrigerated at −80°C for posterior processing.

### CHIKV serology

The sample processing protocol was in accordance with the indications by the commercial house (Abcam^®^ ab177848 anti-CHIKV IgM human ELISA kit and ab177835 anti-CHIKV IgG human ELISA kit). Anti-Ig antibodies were placed on a polystyrene plate. Controls and samples were added to each well, incubated 1 hour at 37°C. A washing was performed and the CHIKV antigen was added to each well and incubated for 20 to 30 minutes at room temperature. Washings were performed, and then biotin-labelled IgG or IgM were added to each well and incubated for 30 minutes at room temperature. Streptavidin conjugated peroxidase (SP), which binds biotinylated antibodies-specific CHIKV to the wells, was washed and added. The TMB substrate was then added which was catalysed by the SP to produce a blue product which changes to yellow for 30 minutes at room temperature and in the dark after the addition of a stop solution. After 15 minutes at room temperature a spectrophotometer read at 450 nm to give the result. The yellow dye density was directly proportional to the amount of IgG or IgM sample as the case for CHIKV captured on the plate.

Analytical specifications according to manufacture states a specificity > 90% and sensibility > 90% for both IgM and IgG anti-CHIKV. No cross reactivity against *Bordetella pertussis, Chlamydia trachomatis, Chlamydia pneumoniae*, Dengue virus, TBE, *Helicobacter pylori*, Herpes Simple Virus 2, Leishmania, Mycoplasma and Schistosoma was found for IgM. No cross reactivity against Dengue virus, Tick born encephalitis, Cytomegalovirus, Epstein Barr virus, and Helicobacter *pylori* was found for IgG.

### Inflammatory biomarkers

High sensitive C-reactive protein (hsCRP) was determined by enhanced turbidimetry with latex particles; reference values positive > 5 mg/dL.

### Statistical analysis

Descriptive analysis was made using means and standard deviation for continuous variables and count and percentages for categorical variables. Two by two tables were used to establish associations between categorical variables. Odds ratios were calculated for associations with 95% confidence intervals using a p value below 5% as statistical significance. A t-student test was used to compare means using also a p value below 5% as statistical significance. Descriptive statistics were depicted in tables and graphics. The Statistical Package for Social Sciences (SPSS) version 22.0 was used for data analysis. GraphPad Prism version 7.04 was used to elaborate figures.

### Ethical considerations

This study was carried out within the ethical standards considered in the declaration of Helsinki 2013. Informed consent was made, prior to the patients’ admission; in that consent was all the information necessary for the patient to make the decision to participate or not in the study. Confidentiality was safeguarded. Case data will not be published or disclosed.

This project is classified within the minimum risk, in accordance with the Resolution No. 008430 of 1993 of the Colombian Ministry of Health, in its Article 11b.

Patients were informed about how to participate in scales, questionnaires, and clinical evaluations, considering the respect for individual privacy and the confidential handling of the results obtained in the study. All information is available for evaluation by approved competent authorities, including all clinical reports that were carried out with patients. The study was approved by the ethics committee from La Universidad de La Sabana (study approval MED-197-2015) and the Hospital Militar Central (study approval 106-2016).

## Results

From the 6528-surveyed people of the COPCORD study, 548 where included in the study because of clinical suspicion of CHIKV infection, as the source of pain. The suspicion was based on patient referral. Some patients were given CHIKV infection diagnosis by a primary care physician upon attendance to a near medical care facility, prior to our evaluation; others simply assumed the diagnosis based on supposed knowledge of the disease, learned from local news media and educational campaigns promoted by the government health organizations.

After applying WHO clinical criteria for CHIKV infection in the 548 patients, only 177 (32.3%) fulfilled the criteria, however when serological confirmation was made (positive values for IgG or IgM), 295 (53.8%) were positive for CHIKV infection. This formed 4 groups of patients: true positives or patients with typical CHIKV symptoms and positive serology (n: 151; 27.6%), true negatives or patients without typical CHIKV symptoms and negative serology (n: 227; 41.4%), false positives or patients with typical CHIKV symptoms without infection (26; 4.6%), and false negatives or patients without typical CHIKV symptoms and confirmed infection (144; 26.3%) (Fig 1). All variables were evaluated according to these four groups.

**Fig 1.**
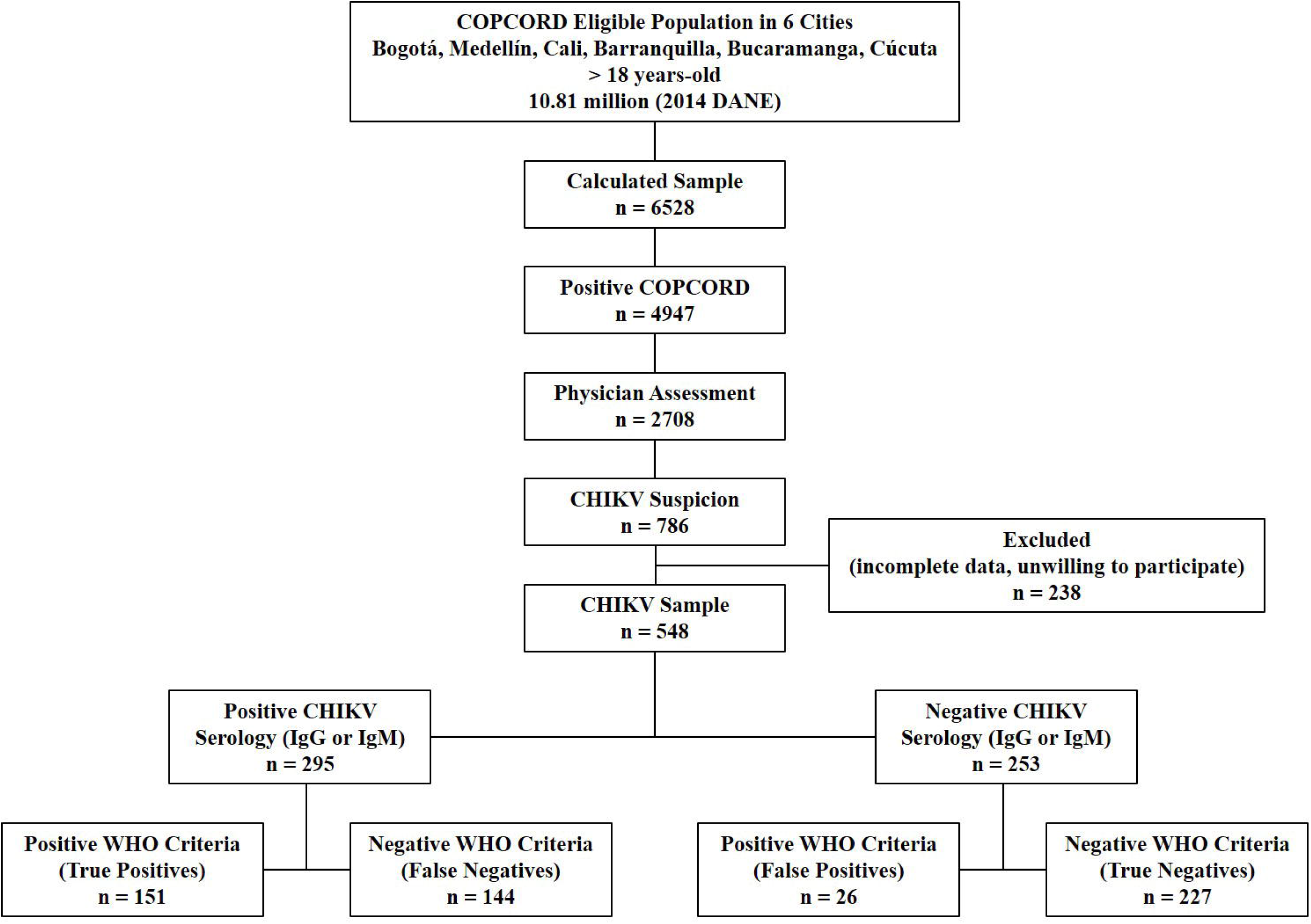
Profile of the study population. DANE: Departamento Administrativo Nacional Estadístico; COPCORD: Community Oriented Program for Control of Rheumatic Diseases; WHO: World Health Organization; CHIKV: Chikungunya Virus

### Demographics

For the complete studied population (548 patients), the mean age was 48.8 years (SD±17.5), where 57.7% (n: 316) were > 45 of age (Table 1). Most of those patients > 45 years (76.3%; n: 241) did not have CHIKV confirmed infection (p = 0.01; OR: 0.63, CI: 0.43-0.93); however most false negatives for CHIKV (66%; n: 95) were in this age group (p = 0.01; OR: 1.60, CI: 1.08-2.38).

**Table 1.**
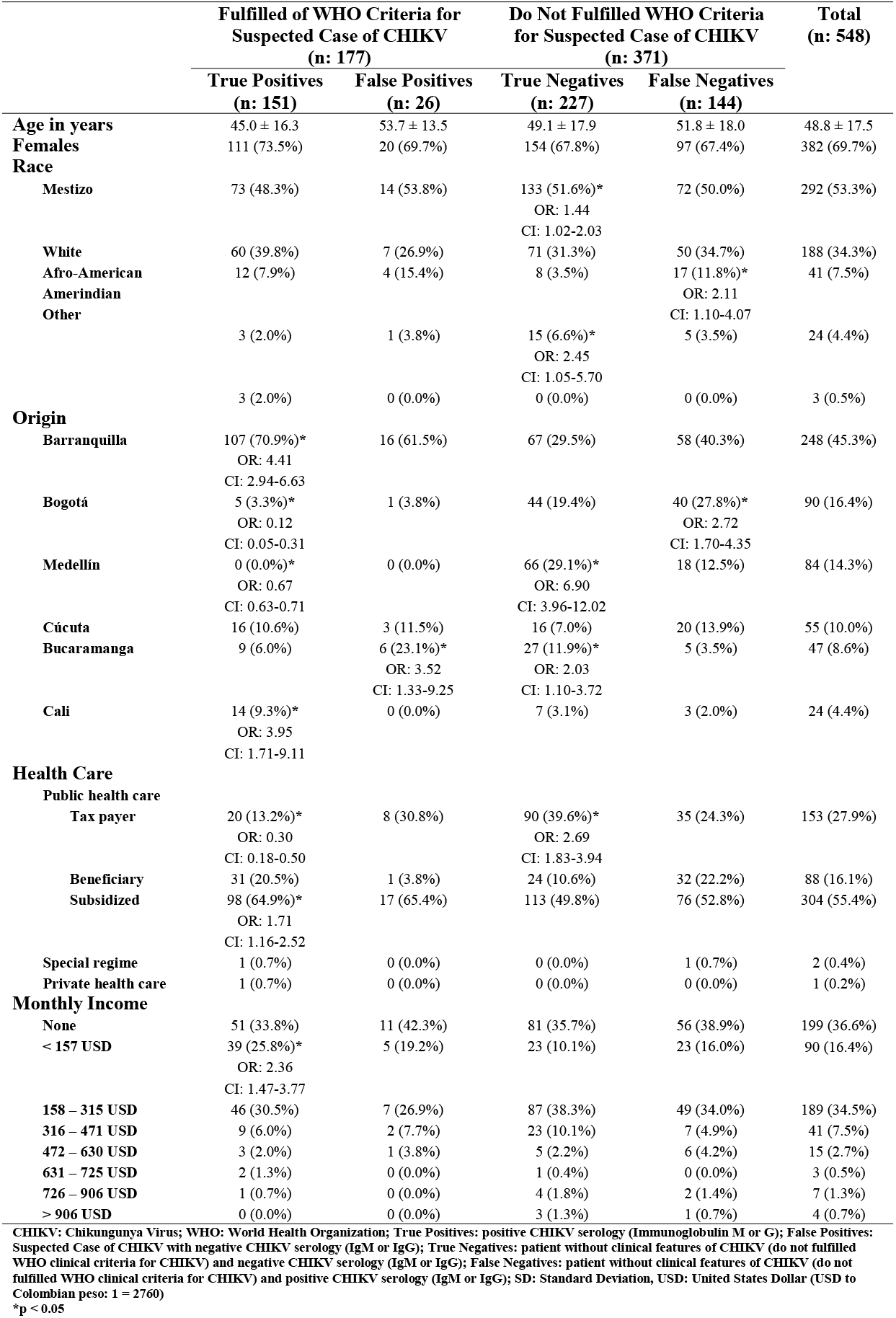
Demographics in patients with suspicion of CHIKV infection

Overall, most patients were female (n: 382, 69.7%) and mestizos (n: 292; 53.35%). Regarding race, most of true negative patients (n: 133; 58.6%) were mestizo (p = 0.03; OR: 1.44, CI: 1.02-2.03), while most Amerindian patients (n: 15; 62.5%) were also true negative (p = 0.03; OR: 2.45, CI: 1.05-5.70). Of interest, there was a higher percentage of false positives (15.4%) and false negatives (11.8%) in Afro-American patients with statistical significance only in the last group (p = 0.02; OR: 2.11, CI: 1.10-4.07) (Table 1).

Barranquilla was the city with most patients in general (n: 248; 45.3%), most of them being true positives (n: 107; 70.9%) conferring a risk of 4.4 times (p = 0.00; OR: 4.41, CI: 2.94-6.63) for confirmed CHIKV infection (Table 1, Fig 2). Another city with increased risk for CHIKV infection was Cali (p = 0.00; OR: 3.95, CI: 1.71-9.11), while living in Bogotá or Medellin was a protective factor (p = 0.00; OR: 0.12, CI: 0.05-0.31 and p = 0.00; OR: 0.67, CI: 0.63-0.71 respectively). Most residents from Medellin (78.6%) and Bucaramanga (57.4%) were true negatives or patients without symptoms and negative serology (p = 0.00; OR: 6.90, CI: 3.96-12.02 and p = 0.02; OR: 2.03, CI: 1.10-3.72 respectively), however, this last city was the second with most false positives (p = 0.07; OR: 3.52, CI: 1.33-9.25). Interestingly, almost half of patients from Bogotá (44.4%) were false negatives or patients without clinical typical symptoms of CHIKV infection but with serologic confirmation of the disease (p = 0.00; OR: 2.72, CI: 1.70-4.35).

**Fig 2.**
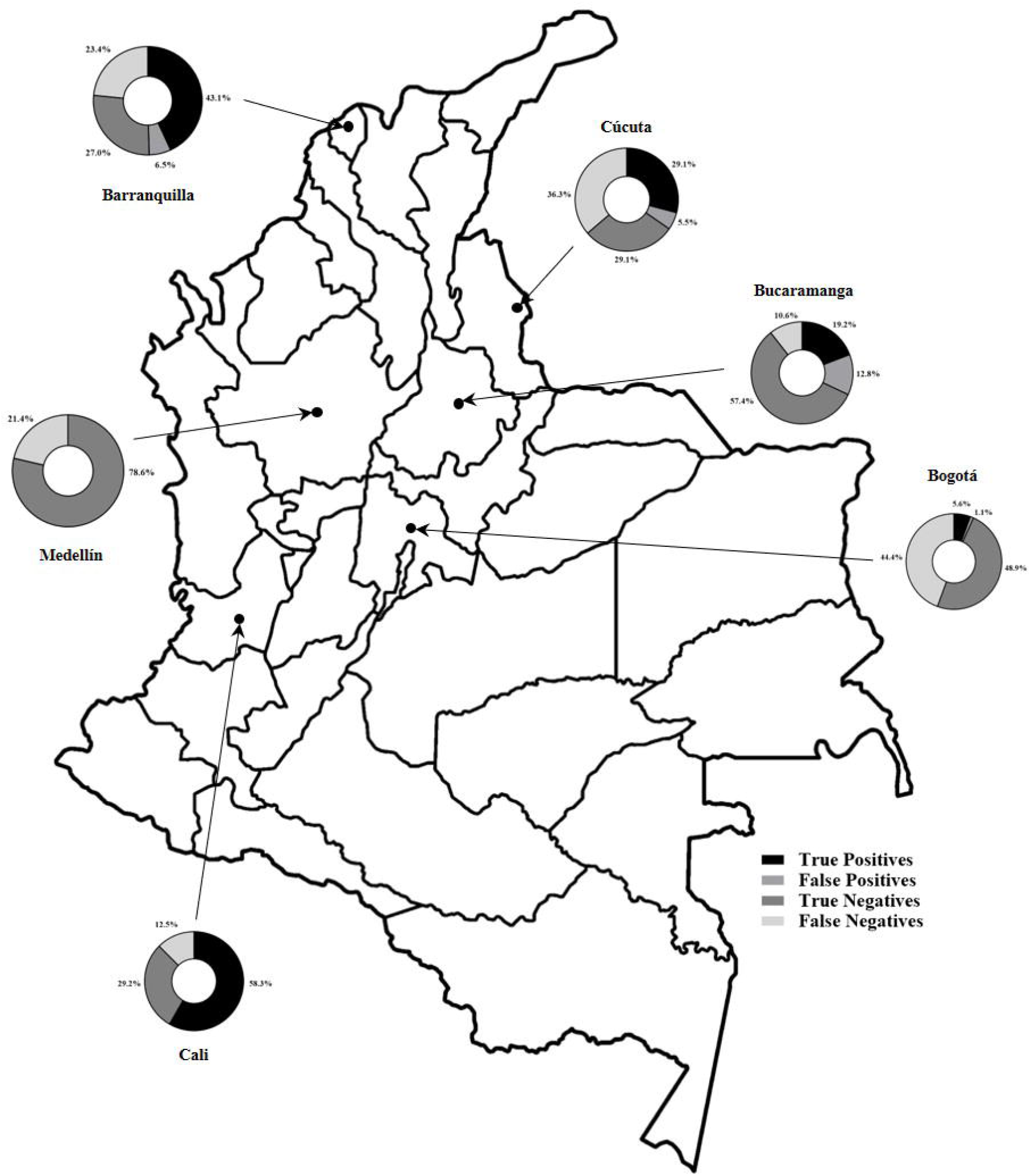
Distribution by city according to WHO criteria and CHIKV serology. WHO: World Health Organization; CHIKV: Chikungunya Virus

In general (S1 Table), many patients lived in a house (n: 422, 77.1%) of their property (n: 412, 75.2%), and most of them were married (n: 164, 29.9%), catholic (n: 412, 75.2%), and had a basic primary school education (n: 182, 33.2%) with knowledge of how to read and write (n: 520, 94.9%). Of interest, 97.4% (n: 221) of true negatives knew how to read and write (p = 0.02; OR: 2.71, CI: 1.08-6.79), while not knowing to read and write was a risk factor for false positives (p = 0.00; OR: 4.13, CI: 1.31-13.02). Also, 19.2% of false negatives patients had no education whatsoever (p = 0.01; OR: 3.31, CI: 1.17-9.31). Almost half of the studied patients (n: 227, 41.4%) were housewives. Of those, 48.3% (n: 73) had confirmation of CHIKV infection (true positives: p = 0.04; OR: 1.47, CI: 1.01-2.15). Only 4 patients with CHIKV confirmed infection (2.6%) were unemployed, which grants a protective factor for being true positive (p = 0.04, OR: 0.34, CI: 0.11-0.99).

Only 0.6% of the studied population had a private health care or special regime (police or military health care) (Table 1). The rest are distributed within different public health care categories which are determined according to the subject’s income. So, people with higher income (tax payer group) must pay an obligatory fee to sustain the other two categories (beneficiary and subsidized) and in this way warrant resources to maintain the health care of those with the lowest income. More than half of patients (55.4%) were in the subsidized health care group, which belongs to lowest income population. Most of these patients (64.9%) were true positive for CHIKV infection granting a risk of 1.7 times (p = 0.00; OR: 1.71, CI: 1.16-2.52), while tax paying patients were protected for the disease (p = 0.00; OR: 0.30, CI: 0.18-0.50) and were almost 3 times more likely to be true negatives (p = 0.00; OR: 2.69, CI: 1.83-3.94).

A hundred and ninety-nine patients (36.6%) had no monthly income, which can be expected considering that most evaluated subjects were housewives. Most of patients with an income below 157 USD per month (n 39, 43.3%) had confirmed CHIKV infection (p = 0.00; OR: 2.36, CI: 1.47-3.77). This socio-economic variable is in relation with a social stratification system implemented by law in Colombia in 1990, where urban populations are classified into different strata with similar economic characteristics (scale from 1 to 6, being 1 the lowest income and 6 the highest) [51]. Overall, 226 analysed subjects were strata 2 (41.2%) followed by strata 1 (n: 169; 30.3%) (S1 Table), representing a majority of low socioeconomic strata. Most of confirmed cases of CHIKV infection patients (n: 72, 47.7%) were strata 1 (p = 0.00; OR: 2.81, CI: 1.90-4.17), while 50.7% (n: 115) of true negatives were strata 2 (p = 0.00; OR: 1.94, CI: 1.37-2.74) and most of strata 4 patients (n: 19, 59.5%) were true negatives (p = 0.03; OR: 1.04-4.47).

### Clinical Features

Of the 548 evaluated patients only 226 recalled duration of symptoms with a mean of 5.53 weeks (±5.92) by the time of the interview. Of those, 53.1% (n: 120) had symptoms for less than 3 weeks (early convalescent), 19.5% (n: 44) between 4 to 6 weeks (late convalescent), and 27.4% (n: 62) for more than 7 weeks (chronic) [52].

Since there were no previous CHIKV infection in Colombia, all patients can be considered as immunologically naïve. Therefore, the serological immunoglobulin (Ig) positivity of each patient can be used as a biological timeline of the infection. So, patients only with positive CHIKV IgM can be considered acute, while patients with positive IgM and IgG can be considered convalescent, and finally patients with only positive IgG can be considered chronic. In our patients, using this classification we found 6.8% (n: 20) of patients with positive IgM (acute), 21.3% (n: 63) with concomitant positivity of IgM and IgG (convalescent), and 71.9% (n: 212) with positive IgG (chronic).

#### Joint Involvement

Eighty six percent of patients suffered from arthralgia (n: 474), with almost 70% being symmetric. In fact, symmetric arthralgia was the most common symptom in all the patients, even when patients are grouped by WHO criteria fulfilment plus CHIKV serology status (as shown in Fig 3). Ninety eight percent (n: 148) of true positive patients referred symmetrical arthralgia (p = 0.00; OR: 34.36, CI: 10.76-104.66), which granted a protective factor for true negative patients (n: 116, 51.1%; p = 0.00; OR: 0.21, CI: 0.14-0.31). However, this feature was also present in false positive (n: 26, 100%; p = 0.00; OR: 1.07, CI: 1.04-1.10), and false negative (n: 92, 63.9%) patients (not statistically significant). This finding could be explained by the population studied, since they are a surrogate from the larger COPCORD sample study, which interviewed subjects that recalled any MSK pain.

**Fig 3.**
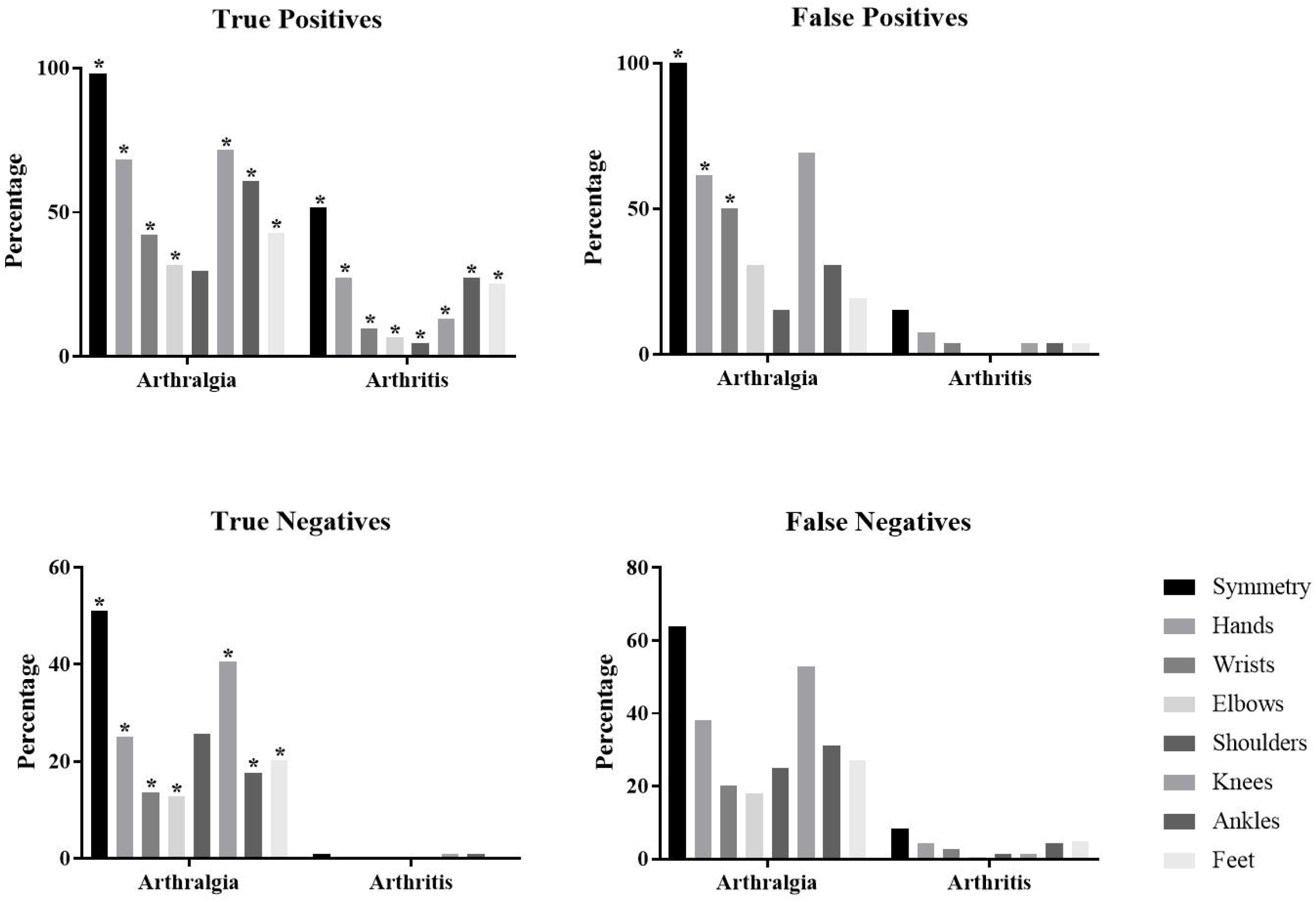
Joint involvement in CHIKV infected patients. CHIKV: Chikungunya Virus; *p < 0.05

Overall, all four groups showed a similar distribution of joint pain involvement, with knees being the most frequent, followed by hands (S2 Table). Of interest, arthralgia in ankles (n: 92, 60.9%; p = 0.00; OR: 3.08, CI: 3.40-7.58) and feet joints (n: 65, 43.0%; p = 0.00; OR: 2.57, CI: 1.73-3.84) had a higher percentage in true positives when compared to false positives, false negatives and true negatives.

A different result was found in true positive patients with symmetric arthritis (n: 78, 51.7%; p = 0.00; OR: 22.49, CI: 12.71-39.80). In general, only 108 patients presented arthritis (19.7%). There was statistically significant difference in arthritis of all the evaluated joints of true positive patients when compared to the other three groups. In true positives, the most frequent swollen joints were ankles (n: 41, 27.2%; p = 0.00; OR: 16.06, CI: 7.57-34.08) and hand joints (n: 41, 27.2%; p = 0.00; OR: 16.12, CI: 8.25-39.79), followed by feet joints (n: 38, 25.2%; p = 0.00; OR: 16.35, CI: 7.41-36.05), knees (n: 20, 13.2%; p = 0.00; OR: 11.96, CI: 4.40-32.52), wrists (n: 15, 9.9%; p = 0.00; OR: 8.64, CI: 3.08-24.23), elbows (n: 10, 6.6%; p = 0.00; OR: 14.00, CI: 3.03-64.70) and shoulders (n: 7, 4.6%; p = 0.00; OR: 9.60, CI: 1.97-46.75).

As mentioned before, to assess the immunologic behaviour of the disease we evaluated the patients’ clinical features according to Ig positivity to CHIKV (Table 2). In general, CHIKV IgM alone was the least frequent Ig found in our studied population (n: 20), whereas IgG was the most present (n: 212). Patients with positive IgG displayed an array of joint symptoms including symmetrical arthralgia and arthritis with statistical significance when compared to the other two groups. Only shoulder pain showed no statistical significance (n: 55; 25.9%). However, some involvement was more common. Knees (n: 141, 66.5%; OR: 2.37, CI: 1.66-3.32), hands (n: 117, 55.2%; OR: 2.39, CI: 1.68-3.41), ankles (n: 105, 49.%; OR: 3.12, CI: 2.16-4.52), feet (n: 76, 35.8%; OR: 1.81, CI: 1.24-2.65) and wrists (66, 31.1%; OR: 1.68, CI; 1.14-2.49) were the most frequent painful joints in IgG positive patients, while ankles (n: 37, 17.5%; OR: 5.25, CI: 2.72-10.14), feet (n: 34, 16.0%; OR: 5.15, CI: 2.60-10.21) and hands (n: 34, 16.0%; OR: 4.08, CI: 2.16-7.71) were the most common swollen joints.

**Table 2.** Joint involvement according to CHIKV Ig positivity

|  | CHIKV Ig Serology |  |  | Total<br>(548) |
| --- | --- | --- | --- | --- |
|  | IgM<br>(20) | IgM + IgG<br>(63) | IgG<br>(212) |  |
| **Arthralgia** |  |  |  |  |
| Symmetry | 11 (55.0%) | 50 (79.4%) | 179 (84.4%)*<br>OR: 3.55<br>CI: 2.31-5.46 | 382 (69.7%) |
| **Hands** | 7 (35.0%) | 34 (14.7%)*<br>OR: 1.71<br>CI: 1.01-2.90 | 117 (55.2%)*<br>OR: 2.39<br>CI: 1.68-3.41 | 231 (42.2%) |
| **Wrists** | 3 (15.0%) | 24 (17.5%)*<br>OR: 2.02<br>CI: 1.16-3.51 | 66 (31.1%)*<br>OR: 1.68<br>CI: 1.14-2.49 | 137 (25.0%) |
| **Elbows** | 2 (10.0%) | 18 (16.2%) | 54 (25.5%)*<br>OR: 1.67<br>CI: 1.09-2.54 | 111 (20.3%) |
| **Shoulders** | 5 (25.0%) | 21 (33.3%) | 55 (25.9%) | 143 (26.2%) |
| **Knees** | 8 (40.0%) | 35 (11.9%) | 141 (66.5%)*<br>OR: 2.37<br>CI: 1.66-3.39 | 294 (53.6%) |
| **Ankles** | 3 (15.0%) | 29 (15.7%)*<br>OR: 1.79<br>CI: 1.05-3.04 | 105 (49.5%)*<br>OR: 3.12<br>CI: 2.16-4.52 | 185 (33.8%) |
| **Feet** | 3 (15.0%) | 25 (16.1%)*<br>OR: 1.79<br>CI: 1.04-3.09 | 76 (35.8%)*<br>OR: 1.81<br>CI: 1.24-2.65 | 393 (71.7%) |
| **Arthritis** |  |  |  |  |
| **Symmetry** | 0 (0.0%)*<br>OR: 0.95<br>CI: 0.93-0.97 | 18 (28.6%)*<br>OR: 2.08<br>CI: 1.14-3.79 | 72 (34.0%)*<br>OR: 6.68<br>CI: 4.04-11.05 | 96 (17.5%) |
| **Hands** | 0 (0.0%) | 13 (20.6%)*<br>OR: 3.24<br>CI: 1.61-6.51 | 34 (16.0%)*<br>OR: 4.08<br>CI: 2.16-7.71 | 49 (8.9%) |
| **Wrists** | 1 (5.0%) | 2 (3.2%) | 16 (7.5%)*<br>OR: 6.77<br>CI: 2.23-20.55 | 20 (3.6%) |
| **Elbows** | 0 (0.0%) | 1 (1.6%) | 10 (4.7%)*<br>OR: 8.26<br>CI: 1.79-38.11 | 12 (2.2%) |
| **Shoulders** | 0 (0.0%) | 2 (3.2%) | 7 (1.3%)*<br>OR: 5.70<br>CI: 1.17-27.71 | 9 (1.6%) |
| **Knees** | 0 (0.0%) | 5 (7.9%) | 17 (8.0%)*<br>OR: 3.57<br>CI: 1.51-8.43 | 25 (4.6%) |
| **Ankles** | 0 (0.0%) | 10 (15.9%) | 37 (17.5%)*<br>OR: 5.25<br>CI: 2.72-10.14 | 50 (9.1%) |
| **Feet** | 0 (0.0%) | 11 (17.5%)*<br>OR: 2.72<br>CI: 1.30-5.67 | 34 (16.0%)*<br>OR: 5.15<br>CI: 2.60-10.21 | 46 (8.4%) |

CHIKV: Chikungunya Virus; Ig: Immunoglobulin, IgM: Immunoglobulin M only; IgG: Immunoglobulin G only
\*p < 0.05

As we use the Ig positivity as a timeline, we observed that initially (in patients with IgM) only 1 patient showed a swollen joint, and few were painful. Later, when IgG starts to appear positive, more joints are involved (painful and swollen), specially hands (n: 13, 20.6%; OR: 3.24, CI: 1.61-6.51) and feet (n: 11, 17.5%; OR: 2.72, CI: 1.30-5.67). By the time IgM fades away, arthritis and arthralgia are present in all the joints. Interestingly, symmetric arthritis confers a protection for IgM positivity in CHIKV suspected patients (OR: 0.95, CI: 0.93-0.97) (Fig 4).

**Fig 4.**
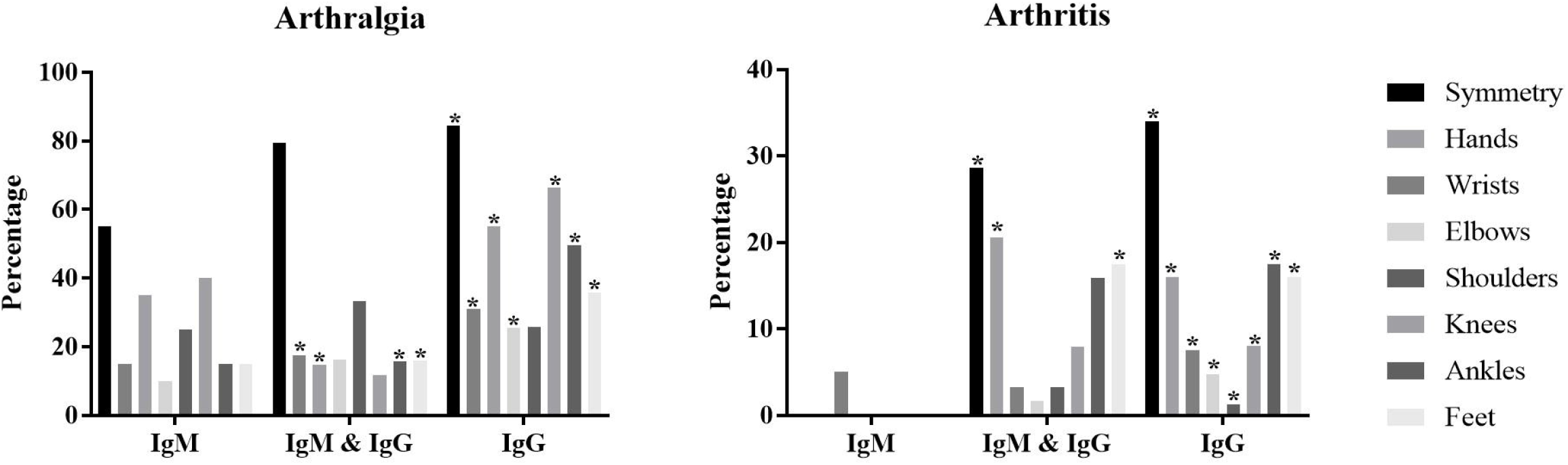
Joint involvement according to CHIKV serology. CHIKV: Chikungunya Virus; IgM: Immunoglobulin M; IgG: Immunoglobulin G; *p < 0.05

We wanted to study the relation between positive hsCRP (> 5 mg/dL) and joint involvement to evaluate active inflammation and clinical features. We found statistical significant relation between positive hsCRP and symmetrical arthralgia (n: 223, 75.3%; p = 0.00; OR: 1.78, CI: 1.23-2.58), arthritis (n: 79, 73.1%; p = 0.00; OR: 2.79, CI: 1.75-4.45) and symmetrical arthritis (n: 72, 75.0%; p = 0.00; OR: 3.05, CI: 1.85-5.02). Regarding specific joint involvement we found that pain in elbows (n: 72, 64.9%; p = 0.01; OR: 1.75, CI: 1.13-2.70), wrists (n: 88, 64.2%; p = 0.00; OR: 1.75, CI: 1.17-2.61), hands (n: 142, 61.5%; p = 0.00; OR: 1.68, CI: 1.19-2.38), ankles (n: 114, 61.6%; p = 0.01; OR: 1.60, CI: 1.11-2.30) and feet (n: 95, 61.3%; p = 0.03; OR: 1.51, CI: 1.03-2.20), as well as arthritis in hands (n: 35, 71.4%; p = 0.01; OR: 2.28, CI: 1.19-4.34), ankles (n: 35, 70.6%; p = 0.01; OR: 2.11, CI: 1.12-3.97) and feet (n: 36, 78.3%; p = 0.00; OR: 3.35, CI: 1.62-6.89), were also statistical significant.

Although we found a relation between clinical presentation (joints involved and type of involvement) and biological timeline markers (presence of IgM, IgM + IgG, and IgG), when we analysed the time of symptoms duration perceived by the patient (< 3 weeks, 4 to 6 weeks and > 7 weeks) and the clinical joint features we found no statistical significance.

#### Systemic Features

To investigate the clinical systemic characteristics of CHIKV infection in our patients we included fever, myalgia and fatigue. These variables were also analysed according to WHO criteria fulfilment (true positives, false positives, true negative, and false negatives), serology positivity (IgM, IgM + IgG, and IgG), hsCRP positivity (> 5 mg/dL) and symptoms’ duration (< 3 weeks, 4 to 6 weeks and > 7 weeks) (Fig 5).

**Fig 5.**
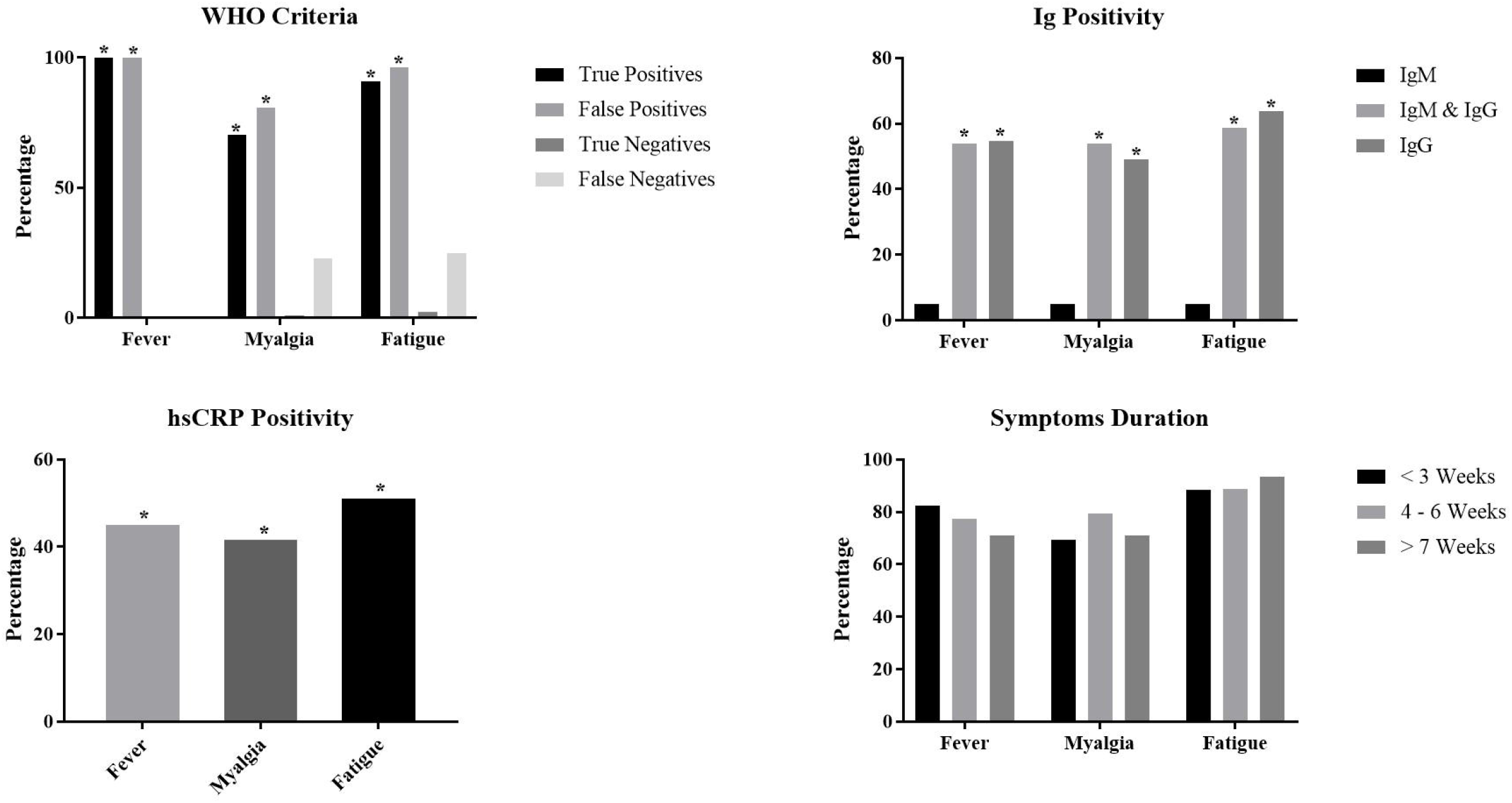
Systemic symptoms in CHIKV patients. CHIKV: Chikungunya Virus; WHO: World Health Organization; IgM: Immunoglobulin M; IgG: Immunoglobulin G; hsCRP: High Sensitive C-Reactive Protein; *p < 0.05

Fever was present in 177 patients in the overall studied population (32.3%) and in all true positive patients (n: 151, 100.0%; p = 0.00; OR: 6.80, CI: 4.77-9.70), however it was also present in patients with negative serology but with symptoms suggestive of CHIKV infection (n: 26, 100.0%; p = 0.00; OR: 1.17, CI: 1.10-1.24). In true negative and false negative patients fever was absent (n: 0, 0.0%; p = 0.00; OR: 0.38, CI: 0.34-0.44 and n: 0, 0.0%; p = 0.00; OR: 0.61, CI: 0.56-0.66 respectively).

Only 1 patient with fever had positive IgM serology for CHIKV (n: 1, 0.5%; p = 0.00; OR: 0.10, CI: 0.01-0.79). Like the pattern observed in joint involvement, more patients suffered from fever as IgG positivity was added to IgM presence (n: 34, 19.2%; p = 0.00; OR: 2.80, CI: 1.64-4.77), and even more so when only IgG persisted (n: 116, 65.5%; p = 0.00; OR: 5.44, CI: 3.69-8.02). As expected, fever was associated with positive hsCRP (n: 135, 76.3%; p = 0.00; OR: 4.19, CI: 2.80-6.27), however this symptom was not associated to symptoms’ duration mentioned by the patient (S3 Table).

Similar findings were obtained for myalgia. The symptom was frequent in true positive patients (n: 106, 70.2%; p = 0.00; OR: 14.34, CI: 9.15-22.77) regardless of most common affected region (generalized myalgia, back myalgia, myalgia in extremities; Table 3). Also, there was a protective factor for CHIKV infection when myalgia was present in true negative patients (n: 2, 0.9%; p = 0.00; OR: 0.009, CI: 0.002-0.037). Again, false positive patients showed high percentage of myalgia (n: 21, 80.8%; p = 0.00; OR: 11.34, CI: 4.19-30.67).

High sensitive CRP positivity as well as presence of IgM and IgG (convalescent) was frequent in patients with myalgia (n: 125, 77.2%; p = 0.00; OR: 4.24, CI: 2.79-6.45 and n: 34, 54.0%; p = 0.00; OR: 3.27, CI: 1.91-5.50 respectively), as well as in patients with positive IgG alone (chronic) (n: 104, 64.2%; p = 0.00; OR: 4.61, CI: 3.12-6.82).

**Table 3.** Myalgia in patients with suspicion of CHIKV infection

|  | **Myalgia** |  |  |
| --- | --- | --- | --- |
|  | **Generalized**<br>**(42)** | **Extremities**<br>**(117)** | **Back**<br>**(26)** |
| **WHO Criteria** |  |  |  |
| **True positives** | 27 (64.3%)*<br>OR: 5.54<br>CI: 2.85-10.76 | 75 (65.0%)*<br>OR: 8.79<br>CI: 5.58-13.85 | 20 (76.9%)*<br>OR: 3.91-25.30<br>CI: 26.66-90.33 |
| **False positives** | 2 (4.8%) | 19 (16.2%)*<br>OR: 11.74<br>CI: 4.80-28.71 | 4 (15.4%)*<br>OR: 4.13<br>CI: 1.31-13.02 |
| **True negatives** | 0 (0.0%)*<br>OR: 0.55<br>CI: 0.51-0.59 | 2 (1.7%)*<br>OR: 0.01<br>CI: 0.004-0.065 | 0 (0.0%)*<br>OR: 0.56<br>CI: 0.52-0.60 |
| **False negatives** | 13 (31.0%) | 20 (17.2%)*<br>OR: 0.51<br>CI: 0.30-0.86 | 2 (7.7%)*<br>OR: 0.22<br>CI: 0.05-0.95 |
| **CHIKV Ig** |  |  |  |
| **IgM alone** | 0 (0.0%) | 1 (0.9%) | 0 (0.0%) |
| **IgM & IgG** | 7 (16.7%) | 27 (23.1%)*<br>OR: 3.29 | 2 (7.7%) |
|  |  | CI: 1.90-5.70 |  |
| **IgG alone** | 33 (78.6%)* | 68 (58.1%)* | 20 (76.9%)* |
|  | OR: 6.69 | OR: 2.76 | OR: 5.72 |
|  | CI: 3.13-14.31 | CI: 1.82-4.20 | CI: 4.69-14.51 |
| **hsCRP** |  |  |  |
| > 5 mg/dL | 32 (76.2%)* | 90 (76.9%)* | 23 (88.5%)* |
|  | OR: 2.93 | OR: 3.64 | OR: 6.99 |
|  | CI: 1.41-6.09 | CI: 2.27-5.82 | CI: 2.07-23.57 |
| **Duration** |  |  |  |
| < 3 weeks | 19 (45.2%) | 62 (53.0%) | 16 (61.5%) |
| 4 to 6 weeks | 11 (26.2%) | 24 (20.5%) | 2 (7.7%) |
| > 7 weeks | 12 (28.6%) | 31 (26.5%) | 8 (30.8%) |

WHO: World Health Organization; Ig: Immunoglobulin IgM: Immunoglobulin M, IgG: Immunoglobulin M, CHIKV: Chikungunya Virus; hsCRP: high sensitivity C Reactive Protein; OR: Odds Ratio; CI: Confidence Interval \*p < 0.05

Regarding fatigue, it was the second most common systemic feature in true positive patients as well as in false positive patients (n: 137, 90.7%; p = 0.00; OR: 49.07, CI: 26.66-90.33 and n: 25, 96.2%; p = 0.00; OR: 48.31, CI: 6.49-359.49 respectively). The symptom was also associated with positive hsCRP (n: 153, 75.4%; p = 0.00; OR: 4.32, CI: 2.94-6.34), and convalescence (n: 37, 58.7%; p = 0.00; OR: 2.73, CI: 1.60-4.67) and chronicity (n: 135, 66.5%; p = 0.00; OR: 6.91, CI: 4.69-10.16) according to CHIKV Ig serology.

Interestingly, we found no relation between the presence of myalgia and fatigue and the duration of symptoms declared by the patient (Table 3).

#### Dermatological Involvement

Skin symptoms were evaluated in all of patients who were suspected of CHIKV infection which corresponded to 27.6% (n: 151). These included the presence of maculopapular rash and site of involvement (face, thorax, abdomen, back, extremities), presence of pruritus, and oral or genital mucosal ulcers.

Maculopapular rash was present in true positive patients (n: 109, 72.2%; p = 0.00; OR: 21.93, CI: 13.59-35.39) as well as in false positive patients although more uncommon in the latter group (n: 14, 53.8%; p = 0.02; OR: 3.27, CI: 1.48-7.26). It was rare in false negatives (n: 23, 16.0%; p = 0.00; OR: 0.41, CI: 0.25-0.67) and even more occasional in true negative patients (n: 5, 2.2%; p = 0.00; OR: 0.02, CI: 0.01-0.06).

In general, in true positive patients the maculopapular rash was more frequent in face, followed by limbs, thorax, abdomen and back, with presence of pruritus in almost half the patients (Table 4). However, it was less frequent in false positive patients with no statistical association regardless of the involved region. Of interest, the presence of maculopapular rash in face and limbs was a protective factor for CHIKV infection in false positive patients (n: 17, 11.8%; p = 0.00; OR: 0.47, CI: 0.27-0.82 and n: 17, 11.8%; p = 0.00; OR: 0.48, CI: 0.27-0.84 respectively).

**Table 4.** Dermatological involvement in patients with suspicion of CHIKV infection

|  | Fulfilled of WHO Criteria for<br>Suspected Case of CHIKV<br>(n: 177) |  | Do Not Fulfilled WHO Criteria<br>for Suspected Case of CHIKV<br>(n: 371) |  | Total<br>(n: 548) |
| --- | --- | --- | --- | --- | --- |
|  | True Positives<br>(n: 151) | False Positives<br>(n: 26) | True Negatives<br>(n: 227) | False Negatives<br>(n: 144) |  |
| Rash |  |  |  |  |  |
| Face | 77 (51.0%)* | 8 (30.8%) | 4 (1.8%)* | 17 (11.8%) | 106 (19.3%) |
|  | OR: 13.20 |  | OR: 0.03 | OR: 0.47 |  |
|  | CI: 8.05-21.65 |  | CI: 0.01-0.10 | CI: 0.27-0.82 |  |
| Thorax | 67 (44.4%)* | 6 (23.1%) | 2 (0.9%)* | 17 (11.8%) | 92 (16.8%) |
|  | OR: 11.86 |  | OR: 0.02 |  |  |
|  | CI: 7.07-19.89 |  | CI: 0.00-0.09 |  |  |
| Abdomen | 67 (44.4%)* | 5 (19.2%) | 2 (0.9%)* | 17 (11.8%) | 91 (16.6%) |
|  | OR: 12.39 |  | OR: 0.02 |  |  |
|  | CI: 7.34-20.91 |  | CI: 0.00-0.09 |  |  |
| Back | 58 (38.4%)* | 5 (19.2%) | 2 (0.9%)* | 15 (10.4%) | 80 (14.6%) |
|  | OR: 10.63 |  | OR: 0.02 |  |  |
|  | CI: 6.19-18.55 |  | CI: 0.00-0.11 |  |  |
| Limbs | 74 (49.0%)* | 10 (38.5%) | 4 (1.8%)* | 17 (11.8%)* | 105 (19.2%) |
|  | OR: 11.34 |  | OR: 0.03 | OR: 0.48 |  |
|  | CI: 6.97-18.44 |  | CI: 0.01-0.10 | CI: 0.27-0.84 |  |
| Mucosal Ulcers |  |  |  |  |  |
| Oral | 7 (4.6%)* | 1 (3.8%) | 0 (0.0%)* | 2 (1.4%) | 10 (1.8%) |
|  | OR: 6.38 |  | OR: 0.57 |  |  |
|  | CI: 1.62-25.02 |  | CI: 0.53-0.62 |  |  |
| **Genital** | 9 (6.0%)* | 1 (3.8%) | 1 (0.4%)* | 2 (1.4%) | 13 (2.4%) |
|  | OR: 6.22 |  | OR: 0.11 |  |  |
|  | CI: 1.88-20.53 |  | CI: 0.01-0.88 |  |  |
| **Pruritus** |  |  |  |  |  |
| **Presence of Pruritus** | 73 (48.3%)* | 12 (46.2%)* | 2 (0.9%)* | 14 (9.7%)* | 101 (18.4%) |
|  | OR: 12.33 | OR: 4.17 | OR: 0.02 | OR: 0.39 |  |
|  | CI: 7.48-20.32 | CI: 1.86-9.31 | CI: 0.00-0.08 | CI: 0.21-0.71 |  |

CHIKV: Chikungunya Virus; WHO: World Health Organization; True Positives: positive CHIKV serology (Immunoglobulin M or G); False Positives: Suspected Case of CHIKV with negative CHIKV serology (IgM or IgG); True Negatives: patient without clinical features of CHIKV (do not fulfilled WHO clinical criteria for CHIKV) and negative CHIKV serology (IgM or IgG); False Negatives: patient without clinical features of CHIKV (do not fulfilled WHO clinical criteria for CHIKV) and positive CHIKV serology (IgM or IgG); OR: Odds Ratio; CI: Confidence Interval
\*p < 0.05

Of the 14 patients with mucosal ulcer involvement, 10 (71.4%) were true positive (p = 0.00; OR: 6.96, CI: 2.15-22.57), 2 (1.4%) were false negative (p= 0.30) and only 1 (7.1%) were false positive (p = 0.66) as well as 1 true negative (p = 0.00; OR: 0.10, CI: 0.01-0.80). Genital ulcers were more frequent in true positive patients (n: 9, 69.2%) with statistical association however oral ulcers were also present in these patients (n: 7; 70.0%). Only few false positive, false negative and true negative patients developed oral or genital ulcers, however only the last group (true negatives) were statistical significant (Table 4).

Pruritus was in almost half the patients with CHIKV suspected patients according to WHO criteria with statistical significance regardless of Ig confirmation, and as expected it was uncommon in true and false negative patients (Table 4).

Regarding skin involvement and CHIKV serology we found few patients with positive IgM (S4 Table). Only 1 patient had pruritic maculopapular rash (5.0%) with no mucosal ulcers. However, along the appearance of positive IgG the number increases to 33 patients with rash (52.4%; p = 0.00; OR: 3.42, CI: 2.00-5.84), 24 with pruritus (38.1%; p = 0.00; OR: 3.26, CI: 1.85-5.73) and 3 with mucosal involvement (4.8%; p = 0.38). The symptoms become even more frequent when only IgG positivity is present being pruritic rash the most frequent skin symptom (46.2%; p = 0.00; OR: 4.59, CI: 3.08-6.83 and 29.2%; p = 0.00; OR: 3.14, CI: 2.01-4.91 respectively). Although there are patients with positive IgG and ulcers they were not enough to achieve statistical significance (n: 9, 4.2%; p = 0.05; OR: 2.93, CI: 0.97-8.88). As mentioned before, few patients had dermatological symptoms when IgM was present however the number increased when IgG was positive. The most frequent areas of involvement, both in patients with IgM plus IgG and IgG positive patients were face and limbs (Fig 6 and S4 Table). Of interest, there was a slight increase number of patients with maculopapular rash in patients with positive IgM plus IgG when compared to patients with IgG alone (Fig 6). Mucosal involvement was rare even in IgG alone patients, nevertheless 9 of 13 patients with genital mucosal ulcers (69.2%) were IgG positive (p = 0.00; OR: 3.68, CI: 1.11-12.10).

**Fig 6.**
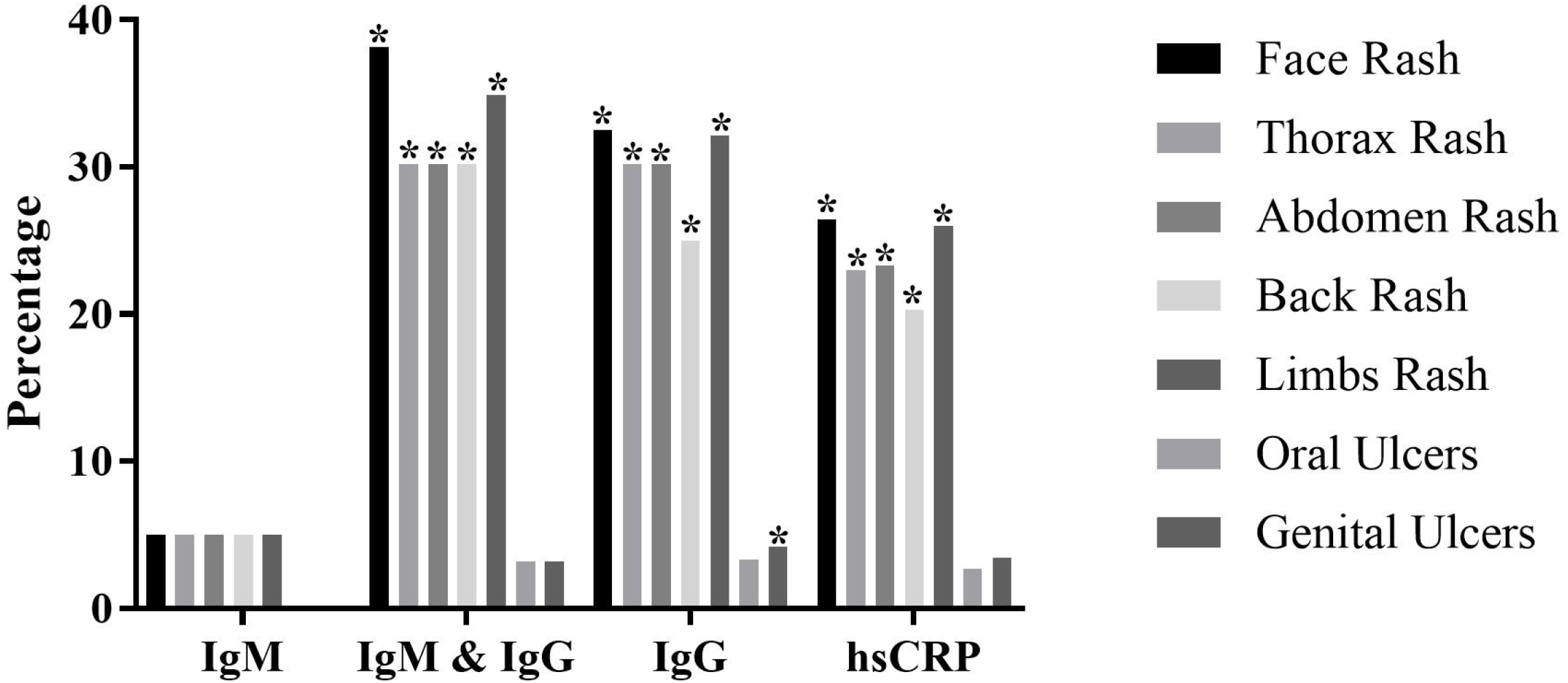
Dermatological symptoms according to CHIKV serology and hsCRP. CHIKV: Chikungunya Virus; IgM: Immunoglobulin M; IgG: Immunoglobulin G; hsCRP: High Sensitive C-Reactive Protein; *p < 0.05

The presence of positive hsCRP was statistically associated with pruritic maculopapular rash in all studied areas but not with mucosal ulcers (S4 Table). There were also no statistical significant associations between symptoms duration and any dermatological involvement in patients with suspicion of CHIKV infection (data not shown).

#### Gastrointestinal Involvement

Overall gastrointestinal symptoms were infrequent in the studied population (n: 103, 18.8%), however in true positive patients were almost in half of the patients (n: 64, 42.4%). For this reason, gastrointestinal involvement was studied with abdominal pain, nausea, emesis and diarrhoea (Table 5).

**Table 5.** Gastrointestinal involvement in patients with suspicion of CHIKV infection

|  | Gastrointestinal Symptoms |  |  |  |
| --- | --- | --- | --- | --- |
|  | Abdominal Pain<br>(30) | Nausea<br>(40) | Emesis<br>(40) | Diarrhoea<br>(90) |
| **WHO Criteria** |  |  |  |  |
| True positives | 12 (7.9%) | 22 (14.6%)<br>OR: 3.59<br>CI: 1.86-6.90 | 18 (11.9%)<br>OR: 2.30<br>CI: 1.20-4.43 | 53 (35.1%)<br>OR: 5.26<br>CI: 3.27-8.46 |
| False positives | 6 (23.1%)<br>OR: 6.22<br>CI: 2.29-16.92 | 5 (19.2%)<br>OR: 3.31<br>CI: 1.17-9.31 | 6 (23.1%)<br>OR: 4.30<br>CI: 1.62-11.43 | 12 (46.2%)<br>OR: 4.87<br>CI: 2.17-10.94 |
| True negatives | 1 (0.4%)<br>OR: 0.04<br>CI: 0.00-0.33 | 1 (0.4%)<br>OR: 0.03<br>CI: 0.00-0.23 | 1 (0.4%)<br>OR: 0.03<br>CI: 0.00-0.23 | 3 (1.3%)<br>OR: 0.03<br>CI: 0.01-0.11 |
| False negatives | 11 (7.6%) | 12 (8.3%) | 15 (10.4%) | 22 (15.3%) |
| **CHIKV Ig** |  |  |  |  |
| IgM alone | 0 (0.0%) | 0 (0.0%) | 0 (0.0%) | 0 (0.0%)<br>OR: 0.95<br>CI: 0.93-0.97 |
| IgM & IgG | 3 (4.8%) | 7 (11.1%) | 5 (7.9%) | 9 (14.3%) |
| IgG alone | 20 (9.4%)<br>OR: 3.39<br>CI: 1.55-7.40 | 27 (12.7%)<br>OR: 3.62<br>CI: 1.82-7.20 | 28 (13.2%)<br>OR: 4.10<br>CI: 2.04-8.27 | 66 (31.1%)<br>OR: 5.87<br>CI: 3.54-9.75 |
| **hsCRP** |  |  |  |  |
| > 5 mg/dL | 22 (7.4%)<br>OR: 2.44<br>CI: 1.07-5.60 | 31 (10.5%)<br>OR: 3.15<br>CI: 1.47-6.76 | 30 (10.1%)<br>OR: 2.72<br>CI: 1.30-5.70 | 67 (22.6%)<br>OR: 2.91<br>CI: 1.75-4.84 |
| **Duration** |  |  |  |  |
| < 3 weeks | 17 (14.2%) | 21 (17.5%) | 20 (16.7%) | 47 (39.2%) |
| 4 to 6 weeks | 5 (11.4%) | 8 (18.2%) | 8 (18.2%) | 16 (36.4%) |
| > 7 weeks | 8 (12.9%) | 11 (17.7%) | 12 (19.4%) | 27 (43.5%) |

WHO: World Health Organization; CHIKV: Chikungunya Virus; hsCRP: high sensitivity C Reactive Protein; OR: Odds Ratio; CI: Confidence Interval \*p < 0.05

Diarrhoea was the most frequent gastrointestinal symptom in true positive patients (n: 53, 35.1%; p = 0.00; OR: 5.26, CI: 3.27-8.46) as well as in false positives (n: 12, 46.2%; p = 0.00; OR: 4.87, CI: 2.17-10.94) and false negative patients (n: 22, 15.3%;), however with no statistical significant association in the last group (p = 0.66). In fact, there were no statistical significant associations in any of the four gastrointestinal symptoms evaluated in false negative patients. Unlike the other 3 groups gastrointestinal involvement was uncommon in true negative patients (Table 5).

In patients with positive IgM, gastrointestinal symptoms were absent (p = 0.02; OR: 0.955, CI: 0.93-0.97) which increased when IgG became positive (n: 13, 20.6%) and IgM fades away (n: 74, 34.9%; p = 0.00; OR: 5.67, CI: 3.53-9.12). Again, diarrhoea was the most frequent gastrointestinal symptom in patients with positive IgM plus IgG (n: 9, 14.3%) and positive IgG alone (n: 66, 31.1%). However only in the last group (positive IgG alone) it reached statistical significant association (p = 0.00; OR: 5.87, CI: 3.54-9.75). In fact, all gastrointestinal symptoms displayed statistical significant association in patients with only positive IgG (Table 5).

Seventy-seven (74.8%) patients out of 103 with positive hsCRP had gastrointestinal symptoms (p = 0.00; OR: 3.05, CI: 1.88-4.94). All four gastrointestinal symptoms studied showed statistical significant association with positive hsCRP, being abdominal pain the less frequent (n: 22, 7.4%; p = 0.02; OR: 2.44, CI: 1.07-5.60) and diarrhoea the most common symptom (n: 67, 22.6%; p = 0.00; OR: 2.91, CI: 1.75-4.84).

Regarding symptoms duration and the presence of gastrointestinal symptoms, we observed a similar distribution within the three studied timeframes (< 3 weeks: 45.8%, 4 to 6 weeks: 40.9% and > 7 weeks: 48.4%) with no statistical significance.

### Disability

To evaluate the disability of the disease at the time of visit, we used three previously validated tools: the health assessment questionnaire disability index (HAQ-DI), the EuroQol-5D (EQ-5D), and an EQ-5D visual analogue scale (VAS). The first two tools were analysed as an absolute number calculating means on groups of patients according to WHO criteria fulfilment, CHIKV Ig presence, hsCRP positivity and symptoms duration according to the patient. Also, HAQ-DI was categorized in mild to moderate disability, moderate to severe disability and severe to very severe disability. The VAS was categorized too in: without disability, mild disability, moderate disability and severe disability. Each category was analysed within the previously mentioned groups.

#### Health Assessment Questionnaire Disability Index (HAQ-DI)

The mean HAQ-DI in the studied population was 0.17 (SD±0.45) (Fig 7). Although there was a similar mean value of HAQ-DI throughout the four groups of patients according to WHO criteria, false positives had a higher value (0.23±0.51) than true positives (0.19±0.45), whereas false negatives had a similar value to true positives (0.18±0.46). Also, patients who referred symptoms for more than 7 weeks had the highest HAQ-DI value (mean: 0.36±0.57). Patients with positive hsCRP, presence of IgG or IgM plus IgG had similar HAQ-DI mean values (0.19, 0.19 and 0.20 respectively), however the lowest value was found in patients with positive IgM (0.05±0.22).

**Fig 7.**
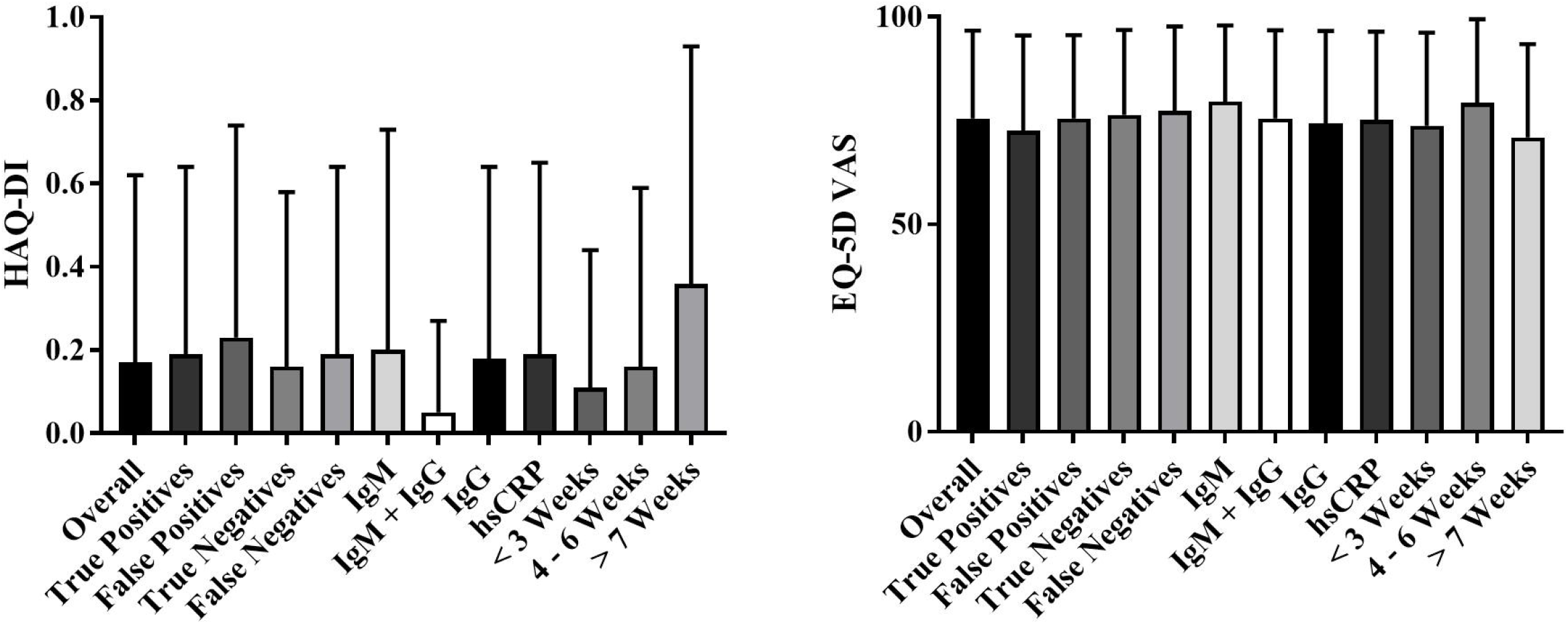
Disability by HAQ-DI and EQ-5D VAS in CHIKV patients. HAQ-DI: Health Assessment Questionnaire Disability Index; EQ-5D: EuroQol-5D; VAS: Visual Analog Scale; CHIKV: Chikungunya Virus; IgM: Immunoglobulin M; IgG: Immunoglobulin G; hsCRP: High Sensitive C-Reactive Protein; *p < 0.05

When HAQ-DI was categorized we found that most patients (95.2 to 100.0%) developed mild to moderate disability regardless of CHIKV serology status, hsCRP positivity, symptoms duration or WHO criteria fulfilment (S5 Table).

#### EuroQol-5D (EQ-5D) Visual Analogue Scale (VAS)

In general, the mean VAS was 75.56 (SD±21.18) from a scale of 0 to 100. We found few differences in VAS values between the studied groups, however false negatives (77.41±20.28), presence of CHIKV IgM (79.65±18.28) and 4 to 6 weeks of symptoms duration (79.30±20.15) had the highest VAS values (Fig 7).

More than half of patients had severe disability (52 to 69.2%) when VAS was categorized (S6 Table). Interestingly, true positive patients were the group with the lowest percentage of severe disability within all studied population (n: 79, 52.3%; p = 0.01; OR: 0.62, CI: 0.42-0.91), while false positives were the ones with highest percentage (n: 18, 69.2%; p = 0.35). Also, around 30 to 40% of patients referred moderate disability according to VAS score regardless of the WHO criteria status, CHIKV Ig presence, hsCRP positivity or symptoms duration. Only 2 patients from the 548 that were studied were without disability by VAS score (0.36%).

#### EuroQol-5D (EQ-5D)

The EQ-5D instrument uses 5 dimensions (mobility, self-care, usual activities, pain/discomfort, anxiety/depression) each one stratified in 3 three categories according to severity, to assess quality of life (Fig 8). Patients were evaluated for associations according to WHO criteria fulfilment, CHIKV Ig serology presence, hsCRP positivity, and symptoms duration.

**Fig 8.**
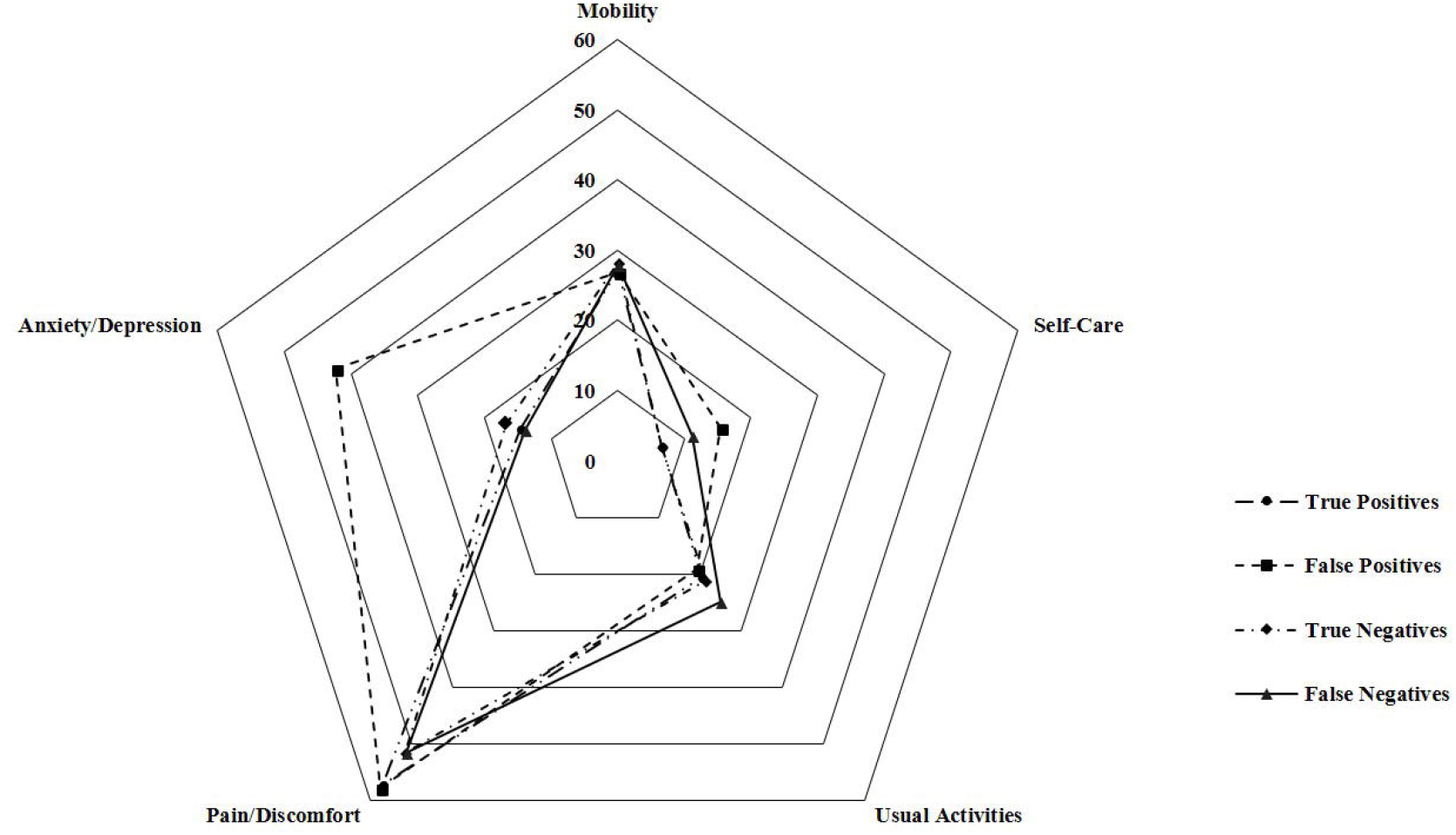
Quality of life according to five dimensions of EQ-5D. EQ-5D: EuroQol-5D

In general, most patients had no problems in the dimensions of mobility, self-care, usual activities and anxiety/depression (72 to 92%), however in the dimension pain/discomfort there was a higher percentage (44.7%) of patients with moderate pain/discomfort (Table 6). In the same way, percentages in all dimensions were similar throughout the four groups according to WHO criteria except for moderate anxiety/depression in false positive patients where the it was higher with statistical significance (n: 10, 38.5%; p = 0.00; OR: 4.17, CI: 1.81-9.57).

**Table 6.** EQ-5D in patients with suspicion of CHIKV infection

|  | Fulfilled of WHO Criteria for<br>Suspected Case of CHIKV<br>(n: 177) |  | Do Not Fulfilled WHO Criteria<br>for Suspected Case of CHIKV<br>(n: 371) |  | Total<br>(n: 548) |
| --- | --- | --- | --- | --- | --- |
|  | True Positives<br>(n: 151) | False Positives<br>(n: 26) | True Negatives<br>(n: 227) | False Negatives<br>(n: 144) |  |
| **Mobility** |  |  |  |  |  |
| No Problems | 110 (72.8%) | 19 (73.1%) | 163 (71.8%) | 104 (72.2%) | 396 (72.3%) |
| Some Problems | 39 (25.8%) | 7 (26.9%) | 63 (27.8%) | 39 (27.1%) | 148 (27.0%) |
| Confined to Bed | 2 (1.4%) | 0 (0.0%) | 1 (0.4%) | 1 (0.7%) | 4 (0.7%) |
| **Self-Care** |  |  |  |  |  |
| No Problems | 141 (93.4%) | 22 (84.6%) | 212 (93.4%) | 128 (88.9%) | 503 (91.8%) |
| Some Problems | 7 (4.6%) | 4 (15.4%) | 15 (6.6%) | 15 (10.4%) | 41 (7.5%) |
| Unable To | 3 (2.0%) | 0 (0.0%) | 0 (0.0%) | 1 (0.7%) | 4 (0.7%) |
| **Usual Activities** |  |  |  |  |  |
| No Problems | 120 (79.5%) | 21 (80.8%) | 179 (78.9%) | 108 (75.0%) | 428 (78.1%) |
| Some Problems | 28 (18.5%) | 5 (19.2%) | 43 (18.9%) | 32 (22.2%) | 108 (19.7%) |
| Unable To | 3 (2.0%) | 0 (0.0%) | 5 (2.2%) | 4 (2.8%) | 12 (2.2%) |
| **Pain/Discomfort** |  |  |  |  |  |
| No | 65 (43.0%) | 11 (42.3%) | 110 (48.5%) | 70 (48.6%) | 256 (46.7%) |
| Moderate | 70 (46.4%) | 12 (46.2%) | 100 (44.1%) | 63 (43.8%) | 245 (44.7%) |
| Extreme | 16 (10.6%) | 3 (11.5%) | 17 (7.4%) | 11 (7.6%) | 47 (8.6%) |
| **Anxiety/Depression** |  |  |  |  |  |
| **No** | 128 (84.8%) | 15 (57.7%)*<br>OR: 0.25<br>CI: 0.11-0.56 | 189 (83.3%) | 124 (86.1%) | 456 (83.2%) |
| **Moderate** | 21 (13.9%) | 10 (38.5%)*<br>OR: 4.17<br>CI: 1.81-9.57 | 29 (12.8%) | 18 (12.5%) | 78 (14.2%) |
| **Extreme** | 1 (0.7%) | 1 (3.8%) | 9 (4.0%) | 2 (1.4%) | 13 (2.4%) |

EQ-5D: EuroQol-5D; CHIKV: Chikungunya Virus; WHO: World Health Organization; True Positives: positive CHIKV serology (Immunoglobulin M or G); False Positives: Suspected Case of CHIKV with negative CHIKV serology (IgM or IgG); True Negatives: patient without clinical features of CHIKV (do not fulfilled WHO clinical criteria for CHIKV) and negative CHIKV serology (IgM or IgG); False Negatives: patient without clinical features of CHIKV (do not fulfilled WHO clinical criteria for CHIKV) and positive CHIKV serology (IgM or IgG); OR: Odds Ratio; CI: Confidence Interval
\*p < 0.05

When EQ-5D were evaluated according to CHIKV Ig presence and hsCRP positivity, most patients had no problems in the mobility (60.0 to 77.8%), self-care (90.0 to 93.7%), and usual activities (53.5 to 81.0%) dimensions, as well as no anxiety/depression (80.0% to 90.5%) (S7 Table). However, there was a higher percentage of moderate pain/discomfort (44.3 to 50.0%) when compared to other dimensions. Specially patients with positive IgM alone had the highest percentage in moderate pain/discomfort, nevertheless it was not statically significant.

Similar findings were found when EQ-5D was evaluated according to symptoms duration. Mobility (67.7 to 77.3%), self-care (86.4 to 95.0%), and usual activities (74.2 to 85.8%) dimensions had higher percentages in the no problem category as well as no anxiety/depression (81.8 to 83.9%) regardless of symptoms duration (S8 Table). Regarding the dimension pain/discomfort it was interesting to find the highest percentage of patients with more than 7 weeks of symptoms in the moderate and extreme category (54.8% and 14.5% respectively). However, patients with symptoms between 4 to 6 weeks had the highest percentage of no pain/discomfort (n: 26, 59.1%; p = 0.00; OR: 2.01, CI: 1.03-3.93).

### Comorbidities

During interview, patients were asked to mention any disease previously diagnosed by a doctor. These diseases were not confirmed by the interviewing doctors; therefore, the information relies on the patient’s memory.

Most patients suffered from smoking (n: 180, 32.8%), followed by hypertension (n: 163, 29.7%), headache (n: 152, 27.7%), peripheral venous insufficiency (n: 126, 23.0%), anxiety and depression (n: 118, 21.5%), obesity (n: 54, 9.9%), diabetes mellitus (n: 48, 8.8%), heart disease (n: 35, 6.4%), cancer (n: 16, 2.9%), stroke (n: 11, 2.0%), epilepsy (n: 7, 1.3%) and tuberculosis (n: 2, 0.4%).

Comorbidities in patients according to WHO criteria for CHIKV infection were distributed like in the general population where smoking was the most frequent comorbidity (Fig 9, S9 Table). However, that was not the case for true positives where only 24.5% were smokers (n: 37; p = 0.01; OR: 0.57, CI: 0.37-0.88) and the most frequent comorbidity was headache (n: 49, 32.5%). Of interest, higher percentages of diabetes mellitus (n: 27, 11.9%; p = 0.02; OR: 1.92, CI: 1.06-3.50) and epilepsy (n: 2, 7.7%; p = 0.00; OR: 8.61, CI: 1.59-46.70) were observed in true negatives and false positives respectively (S9 Table).

**Fig 9.**
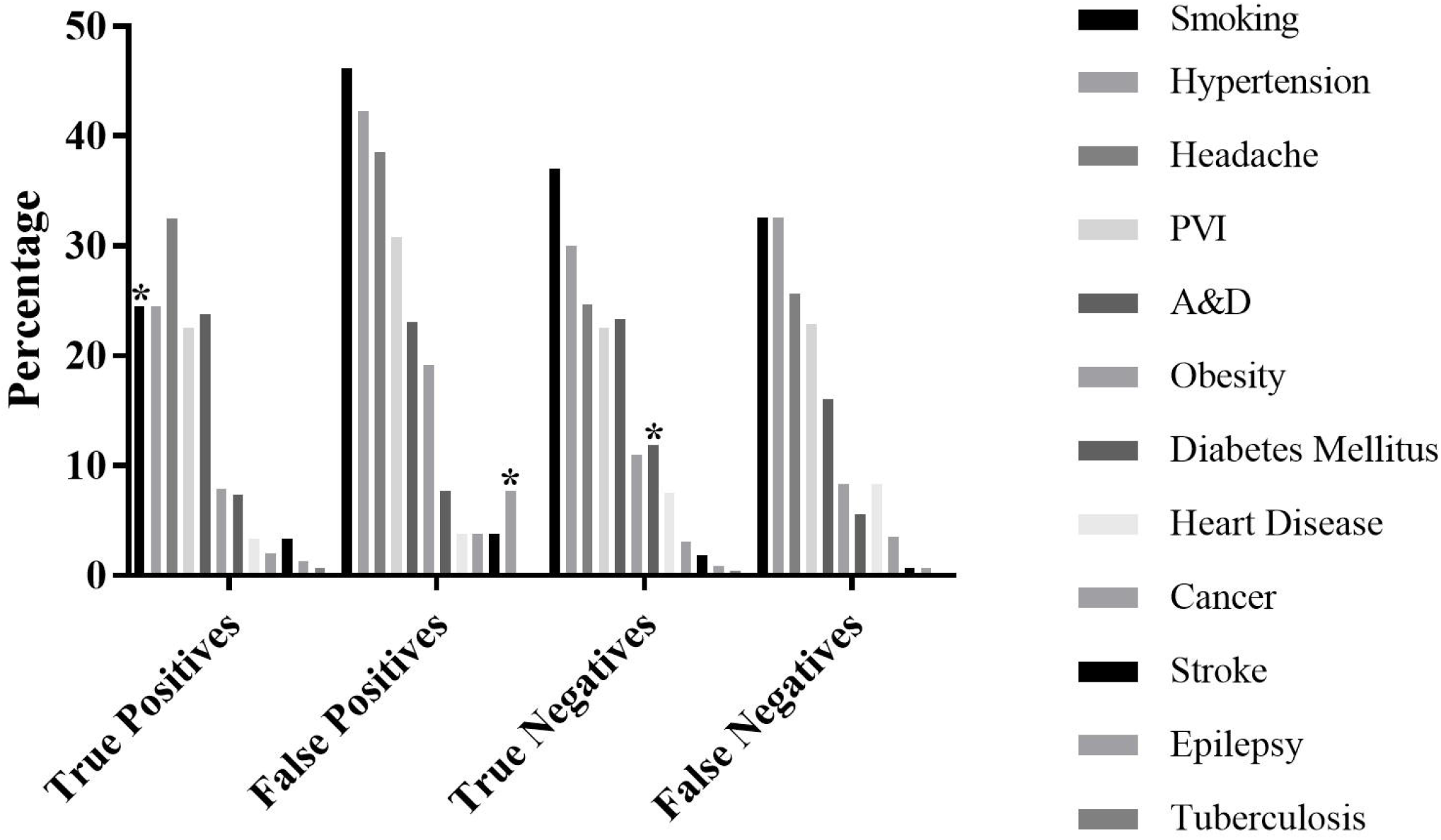
Comorbidities in CHIKV patients. CHIKV: Chikungunya Virus; PVI: peripheral venous insufficiency; A&D: Anxiety and Depression

To evaluate the impact of comorbidity, we decided to analyse their presence according to arthralgia, arthritis, myalgia, fatigue, disability variables (HAQ-DI and EQ-5D VAS) and duration of symptoms (< 3 weeks, 4 to 6 weeks, > 7 weeks) in patients with CHIKV positive serology. Of the 295 patients with CHIKV positive serology only 61 were free of comorbidities.

Overall, arthralgia was the most frequent symptom regardless of associated comorbidity. Only 2 epileptic patients from 3 in total had arthralgia (Table 7). The second most common symptom in CHIKV positive patients with comorbidities was fatigue; patients with stroke had the highest frequency of fatigue (83.3%), followed by cancer (75.0%), diabetes mellitus (73.7%) and headache (67.4%), however only the latter achieved statistical significance (p = 0.04; OR: 1.60, CI: 1.00-2.86). Like in patients without comorbidities, myalgia was the third most common symptom with being the most frequent in patients with cancer (87.5%). Of interest, arthritis was highest in patients with anxiety and depression (52.5%; p = 0.00; OR; 2.73, CI: 1.52-4.90), and less frequent in smokers (23.8%; OR: 0.52, CI: 0.29-0.92).

**Table 7.**
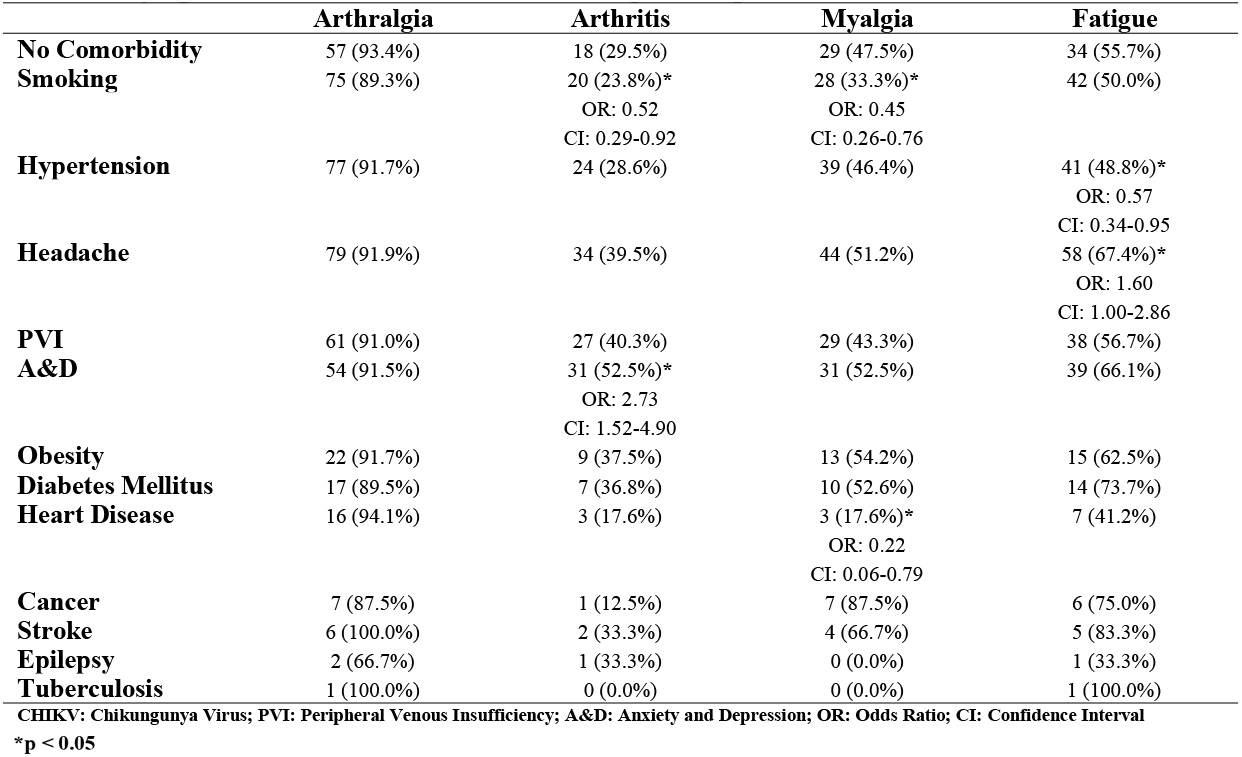
Symptoms and comorbidities in CHIKV positive patients

|  | **Arthralgia** | **Arthritis** | **Myalgia** | **Fatigue** |
| --- | --- | --- | --- | --- |
| **No Comorbidity** | 57 (93.4%) | 18 (29.5%) | 29 (47.5%) | 34 (55.7%) |
| **Smoking** | 75 (89.3%) | 20 (23.8%)*<br>OR: 0.52<br>CI: 0.29-0.92 | 28 (33.3%)*<br>OR: 0.45<br>CI: 0.26-0.76 | 42 (50.0%) |
| **Hypertension** | 77 (91.7%) | 24 (28.6%) | 39 (46.4%) | 41 (48.8%)*<br>OR: 0.57<br>CI: 0.34-0.95 |
| **Headache** | 79 (91.9%) | 34 (39.5%) | 44 (51.2%) | 58 (67.4%)*<br>OR: 1.60<br>CI: 1.00-2.86 |
| **PVI** | 61 (91.0%) | 27 (40.3%) | 29 (43.3%) | 38 (56.7%) |
| **A&D** | 54 (91.5%) | 31 (52.5%)*<br>OR: 2.73<br>CI: 1.52-4.90 | 31 (52.5%) | 39 (66.1%) |
| **Obesity** | 22 (91.7%) | 9 (37.5%) | 13 (54.2%) | 15 (62.5%) |
| **Diabetes Mellitus** | 17 (89.5%) | 7 (36.8%) | 10 (52.6%) | 14 (73.7%) |
| **Heart Disease** | 16 (94.1%) | 3 (17.6%) | 3 (17.6%)*<br>OR: 0.22<br>CI: 0.06-0.79 | 7 (41.2%) |
| **Cancer** | 7 (87.5%) | 1 (12.5%) | 7 (87.5%) | 6 (75.0%) |
| **Stroke** | 6 (100.0%) | 2 (33.3%) | 4 (66.7%) | 5 (83.3%) |
| **Epilepsy** | 2 (66.7%) | 1 (33.3%) | 0 (0.0%) | 1 (33.3%) |
| **Tuberculosis** | 1 (100.0%) | 0 (0.0%) | 0 (0.0%) | 1 (100.0%) |

CHIKV: Chikungunya Virus; PVI: Peripheral Venous Insufficiency; A&D: Anxiety and Depression; OR: Odds Ratio; CI: Confidence Interval

Most patients suffered from mild to moderate disability, like patients without comorbidities (S10 Table). However, the presence of anxiety and depression (p = 0.00; OR: 8.50, CI: 1.52-47.64), stroke (p = 0.01; OR: 11.36, CI: 1.11-115.82) and epilepsy (p = 0.00; OR: 28.70, CI: 2.22-370.55) were associated with moderate to severe disability according to HAQ-DI. Mean HAQ-DI values were higher in patients with stroke (0.83±1.33; p = 0.00), epilepsy (0.67±1.15; p = 0.00) and tuberculosis (0.50±0.71; p = 0.00) when compared to patients without comorbidities (0.05±0.22) (S10 Table and Fig 10).

**Fig 10.**
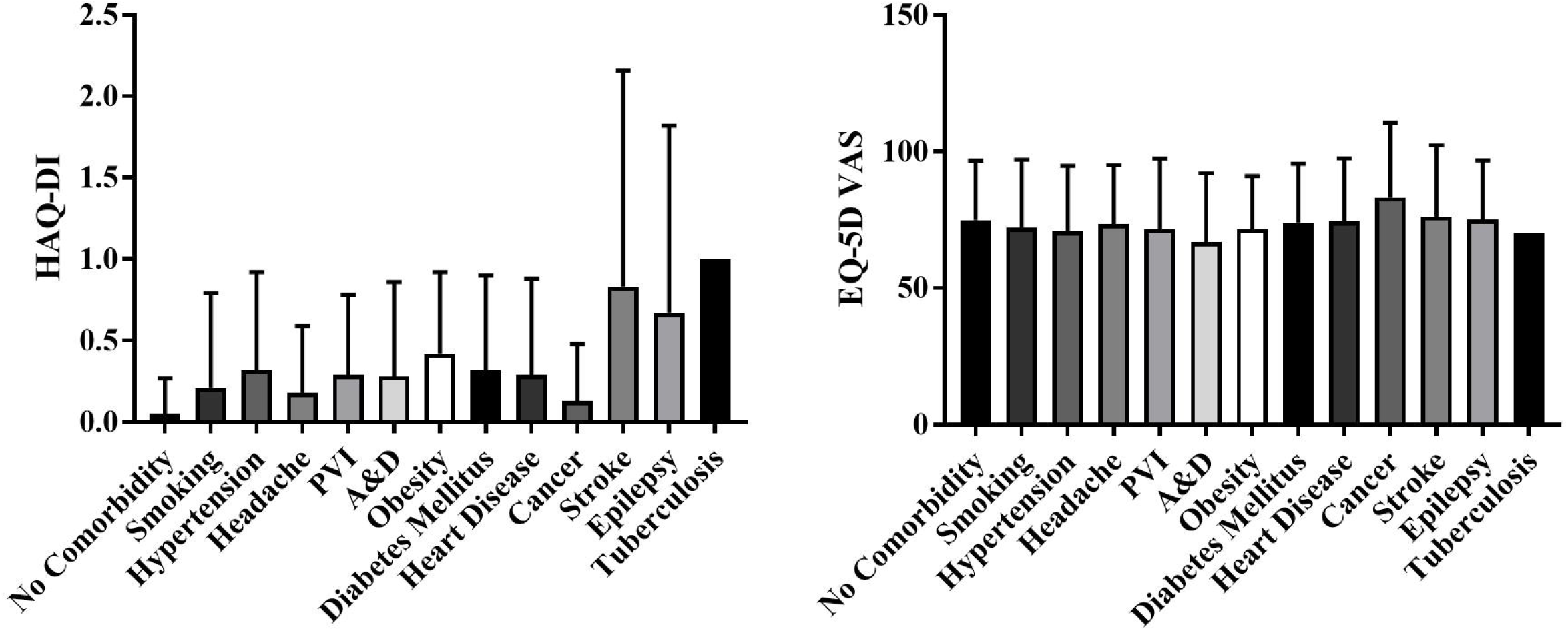
HAQ-DI and EQ-5D VAS of comorbidities in CHIKV patients. HAQ-DI: Health Assessment Questionnaire Disability Index; EQ-5D: EuroQol-5D; VAS: Visual Analog Scale; CHIKV: Chikungunya Virus; PVI: peripheral venous insufficiency; A&D: Anxiety and Depression

Severe disability according to VAS was the most frequent in patients with and without comorbidities (S11 Table). Interestingly more patients with hypertension (p = 0.03; OR: 2.34, CI: 1.03-5.30), peripheral venous insufficiency (p = 0.00; OR: 3.33, CI: 1.46-7.61) and anxiety and depression (p = 0.00; OR: 3.37, CI: 1.46-7.80) suffered from mild disability when compared to patients without a comorbidity. Only patients with cancer referred higher values of VAS (83.13±27.38) when compared to patients without comorbidities (74.90±21.83), though without statistical significance (Fig 10).

Regarding symptoms’ duration, all patients had longer durations regardless of the present comorbidity, where epilepsy (10.50±13.44) and stroke (7.40±7.47) showed the most prolonged periods of symptoms (Table 8). Although most patients had symptoms for less than 3 weeks, there was no statistical significant difference between different categories of symptoms’ duration (Table 8).

**Table 8.** Symptoms’ duration and comorbidities in CHIKV positive patients

|  | Duration |  |  |  |
| --- | --- | --- | --- | --- |
|  | Mean in Weeks | < 3 Weeks | 4 to 6 Weeks | > 7 Weeks |
| **No Comorbidity** | 4.83 (SD±4.94) | 21 (50.0%) | 40 (20.6%) | 55 (28.4%) |
| **Smoking** | 6.21 (SD±6.47) | 23 (48.9%) | 8 (17.0%) | 16 (34.0%) |
| **Hypertension** | 6.17 (SD±5.67) | 20 (41.7%) | 12 (25.0%) | 16 (33.3%) |
| **Headache** | 5.21 (SD±5.54) | 32 (51.6%) | 14 (22.6%) | 16 (25.8%) |
| **PVI** | 6.71 (SD±6.51) | 18 (43.9%) | 7 (17.1%) | 16 (39.0%) |
| **A&D** | 5.43 (SD±5.50) | 23 (54.8%) | 7 (16.7%) | 12 (28.6%) |
| **Obesity** | 5.06 (SD±4.12) | 7 (43.8%) | 4 (25.0%) | 5 (31.3%) |
| **Diabetes Mellitus** | 6.27 (SD±5.64) | 6 (40.0%) | 4 (26.7%) | 5 (33.3%) |
| **Heart Disease** | 5.29 (SD±5.22) | 4 (57.1%) | 0 (0.0%) | 3 (42.9%) |
| **Cancer** | 6.14 (SD±6.49) | 2 (28.6%) | 3 (42.9%) | 2 (28.6%) |
| **Stroke** | 7.40 (SD±7.47) | 1 (20.0%) | 2 (40.0%) | 2 (40.0%) |
| **Epilepsy** | 10.50 (SD±13.44) | 1 (50.0%) | 0 (0.0%) | 1 (50.0%) |
| **Tuberculosis** | 7.0§ | 0 (0.0%) | 0 (0.0%) | 1 (100.0%) |

CHIKV: Chikungunya Virus; PVI: Peripheral Venous Insufficiency; A&D: Anxiety and Depression; SD: Standard Deviation; OR: Odds Ratio; CI: Confidence Interval
\*p < 0.05; § only 1 patient

## Discussion

Colombia is a tropical weather country, located northwest of South America with coasts on the Atlantic and the Pacific Oceans. Since it is a country near the Equator but divided by the Andes mountains, there is a high diversity in altitudes and ecosystems. This allows the habitat of multiple vectors including *Ae aegypti* and *Ae albopictus*, which facilitates the rapid spread of an arthropod-borne disease like CHIKV infection. In 2013 the virus reached the Americas to the Island of Saint Martin [38]. It was a matter of time for CHIKV to reach South American mainland. In Colombia the epidemic started a year later in municipalities near the coast of the Atlantic Ocean [11]. Simultaneously with the CHIKV epidemic, a prevalence study of rheumatic diseases with COPCORD methodology was being carried out. This allowed us to develop a detailed characterization of the CHIKV epidemic in Colombia, with information from multiple regions of the country obtained directly from the patient’s source, decreasing selection bias.

Our study revealed that overall, half of the patients with suspicion of CHIKV infection had the disease via serological confirmation (n: 295, 53.8%). However, it is interesting to find that almost half of those patients did not referred typical symptoms of CHIKV infection (false negatives; n: 144), of which more than a third reside in regions (Bogotá and Medellin) where the CHIKV vector is not endemic (n: 58, 40.3%). This could be explained by the fact that the inhabitants of these regions were not aware of the disease symptoms as well as primary care physicians, resulting in misleading diagnoses and underdiagnosis. The other side of the coin could explain the fact that most false positives were from vector endemic regions (25 out of 26), where the awareness of the disease was higher, leading to overdiagnosis.

Although this theory could be the explanation of some cases, the question arises if there is another reason that could be influencing the lack of CHIKV typical symptoms in these patients, especially the ones from endemic regions (Barranquilla, Bucaramanga, and Cúcuta; 86 of 144). Studies on environmental as well as genetic factors are needed to help elucidate these findings. Also, it is important to note that in our country for the general population and primary care physicians “classical or typical symptoms” of CHIKV infection were the presence of fever plus MSK symptoms. This misconception ends up in discarding patients with MSK symptoms without fever, which in our study, as mentioned before, is one of the causes of false negatives or in the case of the general population an increase in underdiagnosis or underreport. Nevertheless, it exemplifies that in the case of CHIKV infection, a general systemic symptom such as fever has more importance that it should have in diagnostic decision making. Considering that most of endemic viral infections in countries where CHIKV is present also cause fever, the symptom should not be one of the hallmarks for diagnostic criteria.

The mean age of our population study was 48.8 years, however most patients above 45 years old had the infection without fever, or at least without the recall of fever. This can be explained by the fact that most people in this age group are in their labour stage, which leads to pay less attention to any symptom to avoid losing hours or work days.

Like in epidemics from Malaysia, Reunion Island, and Comoro Islands, most studied patients were female, which is expected in a study designed to evaluate patients in their homes during working hours, increasing the probability of assessing housewives [53–56]. A selection bias is present, which must be considered. Nevertheless, to date, it has not been demonstrated that the CHIKV or its vectors have a gender predilection. In fact, a study in Mayotte, found higher seroprevalence in males, suggesting that inconsistencies in gender preponderance are related to differences in exposure due to community-specific habits, customs or behaviours [57].

Our study showed that low income (below 157 USD per month), low socioeconomic strata (strata 1) and poor-quality health care (subsidized health care) increased the risk of CHIKV infection. Although most interviewed subjects had low socio-economic status, which could create a selection bias, this high percentage reflects the current state of the country. These variables are related to poverty, which is consistent with the study in Mayotte where poor living conditions are associated with high risk of CHIKV infection [57].

Another indicator of lower socioeconomic status is education, measured in our study by illiteracy. Although illiteracy was not associated with increased risk of CHIKV infection, it was associated with false positives (patients who suspect having the disease but is not confirmed by serology). We can hypothesize that although most information on CHIKV infection was passed on in the community from person to person, some written campaigns might encourage people with symptoms suggestive of CHIKV infection to attend a primary care physician for a proper diagnosis. Illiterates would not have access to that information, therefore increasing the assumption of the diseases merely based on suspicion, explaining the high percentage of illiterates in false positives. Also, other studies have found that illiteracy is related to less knowledge of mosquito-borne diseases including CHIKV infection, when compared to literate patients, which could confound patients about the real symptoms of the disease as well as decrease awareness of medical consultation [58,59].

One of the biggest problems we encounter during the analysis of the data was the inconsistency in the symptoms timeline between what the patient remembered and the serological biomarkers. While most patients had chronic biomarkers (71.9% were positive only for IgG), only 19.5% of patients recalled symptoms duration for more than 7 weeks. The same discrepancy was found with the presence of acute biomarkers (6.8% with IgM only) and symptoms for less than 3 weeks (53.1%). These findings could explain the lack of significant associations between symptoms duration referred by the patient and clinical symptoms of CHIKV infection, disability and comorbidities. Also, it reveals the great recall bias that our patients showed regarding symptoms duration. Since biomarkers are more accurate, we can conclude that most patients were evaluated during a convalescent-chronic stage of the disease. Nevertheless, we cannot take the findings of our study as a chronic presentation of the disease. For that, we must re-evaluate the patients after two or three years after the first symptoms.

Like other studies, we found symmetrical arthralgia to be a frequent symptom in patients with suspicion of CHIKV infection [30,33,60–62]. Knees, hands and ankles the most affected joints [30,33,60,62–64]. However only arthritis was statistically significant in true positive patients when compared to the other three groups (false positives, false negatives and true negatives). Especially symmetrical arthritis of ankles, hand joints, feet joints and elbows were the sites with higher associations in patients with confirmed CHIKV infection. These findings suggest that symmetric arthritis of the previously mentioned joints is characteristic of CHIKV infection. We believe these symptoms must be cardinal in the diagnostic flow chart of CHIKV infection.

Of interest, we found that the severity of joint symptoms, given by the number of joints involved and the presence of arthritis, is related to the presence of IgG and not IgM. In our patients, as IgG started to appear, more joints were involved both in number and swelling. Likewise, fever, myalgia and fatigue were more present in patients, as CHIKV IgG starts to appear. These findings agree with studies on the immunological response of the host to CHIKV infection. Once the virus reaches the tissues for replication, an innate immune response occurs, which results in production of inflammatory cytokines and chemokines establishing the first response against the infection [65–67]. Especially the production of interleukin 6 (IL-6) induce B lymphocytes and the production of Ig for viral clearance, dominated by IgG [52,68–70]. High levels of IL-6 and IgG are linked to high viremia and the development of symptoms [60,68,70,71].

As expected, we found that fever was a cardinal symptom in CHIKV infected patients. Even more so, when it was almost absent in false positive and true negative patients (patients with negative CHIKV serology). Although this is very compelling, it must be interpreted with care. First, by the time we evaluated the patients the fever was not confirmed by the physical exam. Our study was based on house-to-house interviews and examination, so, some patients were asymptomatic by the time of the physician’s visit. This increase the chance of asymptomatic diagnosis; however, it increases recall bias. Secondly, sometimes the fever was not even confirmed with a thermometer; some patients assumed having fever but did not confirmed it, which could reduce the credibility of the symptom.

Other studies found similar percentages of fever ranging from 90 to 100%, however we have to remember that it is an obligatory symptom for WHO CHIKV infection criteria, increasing selection bias [30,33,75,49,53,54,63,64,72–74]. Another point to consider is that in regions where Zika, Dengue or CHIKV infections can co-exist at the same time, the use of non-specific symptoms like fever in the diagnostic criteria, increases sensibility but decreases specificity reducing the ability to discern which infection is responsible, leading to over or underdiagnosis. An example of the use of more specific symptoms in the diagnosis of an arboviral disease is demonstrated in the study of Braga et al. They found that using the presence of rash, pruritus, conjunctival hyperaemia and excluding fever, anorexia and petechiae increased the performance when compared to other existing Zika suspected case definitions [76]. The same can be extrapolated with other systemic symptoms like fatigue and myalgia (found frequently in our cohort). These are symptoms that although are present in high percentages in CHIKV infection descriptions, can also be found in other arboviral infections [77–79].

Following the same line of overlapping symptoms and arboviral infections is the presence of dermatological involvement. Especially maculopapular rash in CHIKV and Zika infections. Our study showed a higher frequency of maculopapular rash in face, limbs and thorax with pruritus in almost half of the patients. In general, other studies in CHIKV epidemics have showed similar findings [30,80–82]. However, Zika also presents itself with maculopapular rash [77]. In fact, in the previously mentioned study on Zika, maculopapular rash and pruritus is one of the hallmarks of their disease case definition [76]. However, our patients did not refer conjunctivitis and we found an important percentage of mucosal ulcers in CHIKV confirmed patients which are less frequent in Zika and dengue infections. Other studies have found the presence of mucosal ulcers in CHIKV infection, however it appears that it is not frequently evaluated, since it is more mentioned in studies with dermatological focus [80,82,83]. More emphasis should be made in the clinical examination of mucosal ulcers in CHIKV suspected patients, since these symptoms could play an important role in differentiating CHIKV from Zika or dengue infections.

Almost half of the patients with confirmed CHIKV infection suffered from gastrointestinal symptoms, especially diarrhoea. Other studies have found similar frequencies of gastrointestinal involvement; however, it is interesting that these symptoms are not included in the case definition of CHIKV infection [30,49,63,64,73,84]. An explanation may be that other arboviral infections also present gastrointestinal symptoms with some frequency, nevertheless they are far more frequent in CHIKV infection [79,85].

It is well known that CHIKV infection is self-limiting, resolving over time, and with a low mortality rate; however it causes considerable pain and disability, making it a public health issue mainly due to significant economic burden to the health system, families and society [86–89]. A recent study in Colombia estimated an economic cost per CHIKV patient of 152.9 USD, with an overall cost for the epidemic between 2014 and 2015 of 67 million USD [87]. For these reasons we evaluated the disability in our patients using three tools, HAQ-DI, EQ-5D, and EQ-5D VAS.

In general, in our studied population the mean HAQ-DI was low when compared to patients with rheumatic diseases in Colombia (0.49), but higher than the general population (0.01) or patients with non-rheumatic diseases (0.06) [90]. Most patients developed mild to moderate disability, like reports by Bouquillard but with higher HAQ-DI values (0.44) [62]. Rahim et al. also found that most CHIKV patients had mild disability (60%), however they reported a much higher percentage of patients with moderate to severe disability (16%) [91]. The differences between our findings and those of Rahim and Bouquillard’s can be explained by the fact that our population had the infection more recently, while patients from the other two studies were evaluated a year later (15 and 17 months respectively). It is known that patients who develop chronic symptoms will have a higher disability when compared to those who do not [86,88,92,93]. It is probable that an evaluation of our CHIKV positive patients today will show results similar to those mentioned previously.

Most studies on disability in patients with CHIKV infection revealed that depression is commonly present [62,92,94–96]. In fact, some theories have been postulated to explain this finding, where the high levels of IL-6 present in CHIKV infected patients can be related to depression since this cytokine is also elevated in depressed patients [97–99]. However, we did not find high percentages in the anxiety/depression dimension. Again, this could be explained by the time our patients were studied during the course of the disease. We believe that higher percentages of depression in CHIKV positive patients is a result of chronification of the disease.

As expected, patients with comorbidities, especially those that are more disabling like stroke or epilepsy, had higher values of HAQ-DI and EQ-5D VAS. Nevertheless, the were no statistically significant differences in severity between patients with and without comorbidities measured by symptoms or duration of symptoms. In this regard, we found that only 61 patients of the 295 with positive CHIKV were free of comorbidities. In the remaining 234 patients, the most frequent comorbidity was hypertension, followed by headache and peripheral venous insufficiency, which is similar to that found in patients with rheumatic diseases (Londono et al. article under review).

Our study has some limitations. First, there is a selection bias regarding MSK symptoms since the study was developed within the framework of a COPCORD methodology. Also, another selection bias is related to the group of subjects evaluated in a house-to-house system; in our case women housewives. However, the fact that the patients were evaluated in their homes and not on a medical setting decreases the chance of capturing more severe cases. This allows to find even asymptomatic patients that otherwise would not attend a physician. Also, makes it easier to get a more real idea of the epidemic. Second, there is always the recall bias of the symptoms that could not be corroborated by the medical evaluator. Third, there were no confirmation of the disease by protein chain reaction due to high costs. However, considering that in Colombia there has never been a chikungunya epidemic, it is assumed that the population is immunologically naïve, therefore we did not see the need to confirm the diagnosis. Also, since some patients were evaluated after the infection, we were not able to do viral loads which would have been helpful to create an adequate timeline of the infection. Lastly, we did not perform dengue serology to evaluate cross reactions with CHIKV, however it is important to note that the commercial house of the used kits ensure no cross reactivity and specificity and sensibility above 90% for both IgG and IgM.

On the other hand, our study has some strengths. One, since it was developed within the framework of a COPCORD methodology, a great amount of data could be gathered and compared with healthy population and rheumatic disease patients. Also, a reasonable representation of the country’s population is represented in the study, considering the number of evaluated patients and the six completely different cities where the study took place. Another strength is the fact that all patients were evaluated by a rheumatologist or a rheumatology fellow which warrant the accuracy of physical examination, especially the musculoskeletal system. Finally, the long period in which the study was carried out allowed the collection of data during different moments of the epidemic.

We believe that our study describes a global, complete and detailed view of the CHIKV epidemic in Colombia, that not only leaves a ground base to develop future research that will hopefully answer the questions raised during its progress, but also raise concern on the burden and impact of the disease in the Colombian population, that can be used by governmental entities.

## Conclusions

Our study showed that poverty and low socioeconomic status are associated with increased risk of CHIKV infection. Public health strategies on prevention, education and vector control should prioritise vulnerable communities as well as decreasing health inequalities and social disparities. There is a considerable number of patients that do not display the “typical or classical” symptoms of CHIKV infection which leads to false negatives, underreport and underdiagnosis. Studies are needed to investigate the reasons why some patients display fever and other do not as well as why MSK are more severe or chronic in some patients than in others.

A distinctive clinical picture is presented by CHIKV infection with symmetrical arthritis of hand joints, ankles, and feet joints as the hallmark for diagnostic clinical criteria. Although fever, myalgia and fatigue are present in high percentages in CHIKV confirmed patients, Zika and dengue infections can also produce these symptoms which decreases their ability to correctly make diagnose from clinical stand point in a primary care setting. More distinctive clinical features of each disease are more useful for the primary care physician to diagnose arboviral diseases.

Data recollection in epidemics is bound to recall bias by the patient, which could cause misleading results. This is an issue that always must be considered, however the use of biomarkers helps reduce bias and allows for accurate clinical associations.

Although not as representative as in rheumatic patients, acute CHIKV causes disability that eventually is represented in economic burden.

## Acknowledgments

The study was financed by the Colombian Rheumatology Association (ASOREUMA Acta 169 10^th^ July 2015), grants from the Universidad de La Sabana (grant number MED-197-2015), Hospital Militar Central (grant number 106-2016) and COLCIENCIAS doctoral scholarship (grant number 757-2016).

The authors would like to thank the EuroQol Research Foundation for allowing a partnership with La Universidad de La Sabana which allowed the use of the EQ-5D instrument.

## Conflict of interest

The authors declare no conflict of interest.

## Supporting information

**S1 Table. Demographics in patients with suspicion of CHIKV infection**. CHIKV: Chikungunya Virus; WHO: World Health Organization; True Positives: positive CHIKV serology (Immunoglobulin M or G); False Positives: Suspected Case of CHIKV with negative CHIKV serology (IgM or IgG); True Negatives: patient without clinical features of CHIKV (do not fulfilled WHO clinical criteria for CHIKV) and negative CHIKV serology (IgM or IgG); False Negatives: patient without clinical features of CHIKV (do not fulfilled WHO clinical criteria for CHIKV) and positive CHIKV serology (IgM or IgG); *p < 0.05

**S2 Table. Joint involvement in patients with suspicion of CHIKV infection**. CHIKV: Chikungunya Virus; WHO: World Health Organization; True Positives: positive CHIKV serology (Immunoglobulin M or G); False Positives: Suspected Case of CHIKV with negative CHIKV serology (IgM or IgG); True Negatives: patient without clinical features of CHIKV (do not fulfilled WHO clinical criteria for CHIKV) and negative CHIKV serology (IgM or IgG); False Negatives: patient without clinical features of CHIKV (do not fulfilled WHO clinical criteria for CHIKV) and positive CHIKV serology (IgM or IgG); OR: Odds Ratio; CI: Confidence Interval; *p < 0.05

**S3 Table. Systemic features in patients with suspicion of CHIKV infection**. WHO: World Health Organization; CHIKV: Chikungunya Virus; hsCRP: High Sensitivity C Reactive Protein; OR: Odds Ratio; CI: Confidence Interval; *p < 0.05

**S4 Table. Skin involvement according to CHIKV Ig and hsCRP positivity**. CHIKV: Chikungunya Virus; Ig: Immunoglobulin, hsCRP: high sensitivity C Reactive Protein; IgM: Immunoglobulin M only; IgG: Immunoglobulin G only; OR: Odds Ratio; CI: Confidence Interval; *p < 0.05

**S5 Table. HAQ-DI in patients with suspicion of CHIKV infection**. HAQ-DI: Health Assessment Questionnaire Disability Index; WHO: World Health Organization; Ig: Immunoglobulin IgM: Immunoglobulin M, IgG: Immunoglobulin M, CHIKV: Chikungunya Virus; hsCRP: high sensitivity C Reactive Protein

**S6 Table. EQ-5D VAS in patients with suspicion of CHIKV infection**. EQ-5D: EuroQol-5D; VAS: Visual Analogue Scale; WHO: World Health Organization; Ig: Immunoglobulin IgM: Immunoglobulin M, IgG: Immunoglobulin M, CHIKV: Chikungunya Virus; hsCRP: high sensitivity C Reactive Protein; OR: Odds Ratio; CI: Confidence Interval; *p < 0.05

**S7 Table. EQ-5D according to CHIKV Ig and hsCRP positivity**. EQ-5D: EuroQol-5D; CHIKV: Chikungunya Virus; Ig: Immunoglobulin, hsCRP: high sensitivity C Reactive Protein; IgM: Immunoglobulin M only; IgG: Immunoglobulin G only

**S8 Table. EQ-5D and duration of symptoms**. EQ-5D: EuroQol-5D; OR: Odds Ratio; CI: Confidence Interval; *p < 0.05

**S9 Table. Comorbidities in patients with suspicion of CHIKV infection**. CHIKV: Chikungunya Virus; WHO: World Health Organization; True Positives: positive CHIKV serology (Immunoglobulin M or G); False Positives: Suspected Case of CHIKV with negative CHIKV serology (IgM or IgG); True Negatives: patient without clinical features of CHIKV (do not fulfilled WHO clinical criteria for CHIKV) and negative CHIKV serology (IgM or IgG); False Negatives: patient without clinical features of CHIKV (do not fulfilled WHO clinical criteria for CHIKV) and positive CHIKV serology (IgM or IgG); PVI: Peripheral Venous Insufficiency; A&D: Anxiety and Depression OR: Odds Ratio; CI: Confidence Interval; *p < 0.05

**S10 Table. HAQ-DI in CHIKV positive patients with comorbidities**. HAQ-DI: Health Assessment Questionnaire Disability Index; VAS: Visual Analogue Scale; PVI: Peripheral Venous Insufficiency; A&D: Anxiety and Depression; OR: Odds Ratio; CI: Confidence Interval; *p < 0.05; § only 1 patient

**S11 Table. EQ-5D VAS in patients with comorbidities**. EQ-5D: EuroQol-5D; VAS: Visual Analogue Scale; PVI: Peripheral Venous Insufficiency; A&D: Anxiety and Depression; OR: Odds Ratio; CI: Confidence Interval; *p < 0.05

